# Protein complexes in cells by AI-assisted structural proteomics

**DOI:** 10.1101/2022.07.26.501605

**Authors:** Francis J. O‘Reilly, Andrea Graziadei, Christian Forbrig, Rica Bremenkamp, Kristine Charles, Swantje Lenz, Christoph Elfmann, Lutz Fischer, Jörg Stülke, Juri Rappsilber

## Abstract

Accurately modeling the structures of proteins and their complexes using artificial intelligence is revolutionizing molecular biology. Experimental data enables a candidate-based approach to systematically model novel protein assemblies. Here, we use a combination of in-cell crosslinking mass spectrometry, cofractionation mass spectrometry (CoFrac-MS) to identify protein-protein interactions in the model Gram-positive bacterium *Bacillus subtilis*. We show that crosslinking interactions prior to cell lysis reveals protein interactions that are often lost upon cell lysis. We predict the structures of these protein interactions and others in the *Subti*Wiki database with AlphaFold-Multimer and, after controlling for the false-positive rate of the predictions, we propose novel structural models of 153 dimeric and 14 trimeric protein assemblies. Crosslinking MS data independently validates the AlphaFold predictions and scoring. We report and validate novel interactors of central cellular machineries that include the ribosome, RNA polymerase and pyruvate dehydrogenase, assigning function to several uncharacterized proteins. Our approach uncovers protein-protein interactions inside intact cells, provides structural insight into their interaction interface, and is applicable to genetically intractable organisms, including pathogenic bacteria.

## Introduction

Life depends on functional interactions between biological macromolecules, with those between proteins being the most diverse and numerous. The structure of protein-protein interactions (PPIs) is inextricably linked to their function, and elucidating these structures is normally laborious. Both proteomic and genetic approaches have been used to compile vast lists of protein-protein interactions, but provide little insight into the topology of the proposed PPIs. Although proteome-wide PPI modeling has been attempted by relying on docking algorithms driven by evolutionary contacts (Cong *et al*, 2019; Green *et al*, 2021), these are limited when detecting dramatic conformational changes upon binding. The recent development of AlphaFold-Multimer brought accurate predictions of the structure of protein-protein complexes into reach (Evans *et al*, 2022; Mirdita *et al*, 2022). This makes establishing structure-function relationships across whole interactomes a possibility (Burke *et al*, 2021; Hopf *et al*, 2014; Akdel *et al*, 2022), and offers a plausible remedy to the Understudied Proteins Challenge (Kustatscher *et al*, 2022), opening a new era in structural systems biology.

There are large caveats for applying AlphaFold-Multimer to model protein interactions across proteomes, however. Predicting the interaction interfaces of all possible combinations of protein pairs is prohibitively expensive and computationally impractical. For example, the 4,257 protein coding genes in *Bacillus subtilis* (Borriss *et al*, 2018) result theoretically in 9 million pairs and 38 billion trimers. While this is already a computational challenge, proteins also form complexes involving much larger numbers of subunits. It has thus become of interest to find shortcuts towards identifying the topology of these interactions, ideally without laborious experimental approaches.

Large numbers of PPIs have been experimentally identified by two-hybrid, affinity purification mass spectrometry (AP-MS) and cofractionation MS studies (CoFrac-MS), among others, for many biological systems from bacteria and yeast to human cells (Gavin *et al*, 2006; Wan *et al*, 2015; Rajagopala *et al*, 2014; Iacobucci *et al*, 2021; Fossati *et al*, 2021). These techniques report thousands of interactions with varying accuracy. However, they provide little topological or structural information, often even leaving open if an interaction is direct or indirect. Additionally, they involve probing interactions outside their native environment, either by lysing the cell or by creating fusion constructs. These approaches therefore systematically miss weak interactions that are lost upon cell lysis and interactions that depend on the native environment of the cell. This may be a substantial contributor to the Understudied Proteins Challenge that sees many proteins currently left without any known function (Kustatscher *et al*, 2022). Nevertheless, the current information in PPI databases has been used as a basis for AlphaFold protein interaction screens in *Escherichia coli (Gao et al, 2022), Saccharomyces cerevisiae* (Humphreys *et al*, 2021) and human proteomes (Burke *et al*, 2021). Unfortunately, it is unknown how many false positive or false negative protein interaction structure predictions this produces. A possible solution would be provided by in-cell structural data that can feed into and independently validate protein structure predictions at scale.

In recent years, *in vivo* crosslinking of proteins and subsequent identification of the linked residue pairs by mass spectrometry (crosslinking MS) has emerged as a technique that can detect PPIs in cells and provide topological information on these interactions (O‘Reilly *et al*, 2020; Chavez *et al*, 2018), with tightly controlled error rates (Lenz *et al*, 2021). By fixing interactions inside cells as the first step of the analytical workflow and providing information on the linked residue pairs, it provides insights into the structure of protein-protein interactions in their native context. However, it remains to be shown whether this may systematically capture interactions that are easily lost upon cell lysis.

Here we combine crosslinking MS and CoFrac-MS of crosslinked cells, two complementary experimental in-cell PPI mapping approaches, to discover PPIs in the Gram-positive model bacterium *B. subtilis* and prove that crosslinking indeed captures otherwise elusive interactions within cells. *B. subtilis* is a major workhorse for commercial protein production and a close relative to the human pathogens *Bacillus anthracis, Listeria monocytogenes* and *Staphylococcus aureus* (Errington & Aart, 2020; Kovács, 2019). Despite its importance as a model organism for Gram-positive bacteria, no systematic PPI screen has been performed in *B. subtilis* so far. Thus, annotation of its PPIs relies on genetic data, targeted biochemical experiments and homology to those reported (from high-throughput screens) in other species. Crosslinking MS provides information on PPI topology, but currently lacks depth in the context of whole-proteome analyses. In contrast, CoFrac-MS can infer the subunits of soluble complexes, but does not provide topological information (Skinnider & Foster, 2021).

To generate structural models of interactions across the *B. subtilis* proteome, we submitted our experimentally-derived PPIs and previously annotated interactions found in the *Subti*Wiki database (Pedreira *et al*, 2022) to protein structure modeling using AlphaFold-Multimer. Importantly, we used a target-decoy approach to benchmark the predicted interface TM-score (ipTM) (Zhang & Skolnick, 2007; Evans *et al*, 2022) in this study. Using the stringent cut-off ipTM > 0.85, we predicted first high-quality structural models for 130 binary protein assemblies, 17 of which are novel in both association and structure. The pairwise interactions can be used as building blocks for further structure predictions of novel higher-order complexes. With this approach we identify the previously uncharacterized protein YneR, here renamed PdhI, as an inhibitor of the pyruvate dehydrogenase, which links glycolysis and the Krebs cycle. In this case, experimental data from global proteomic approaches, structure modeling, and *in vivo* validation converge to identify a novel protein-protein interaction and to demonstrate its biological function. This workflow demonstrates the power of combining complementary techniques including in-cell crosslinking to discover high-confidence direct protein interactions without genetic modification, and to accurately predict and validate corresponding structural models.

## Results

### Crosslinking MS to identify protein-protein interactions within intact *B. subtilis* cells

We generated a whole cell interaction network using crosslinking mass spectrometry. We crosslinked proteins in *B. subtilis* cells with the membrane permeable crosslinker DSSO (Kao *et al*, 2011; Kolbowski *et al*, 2022). Cells were lysed, the proteins fractionated and trypsin digested, and the resulting peptides separated by cation exchange and size exclusion chromatography prior to mass spectrometry and database searching to result in three datasets (**Fig. S1A** and methods). A 2% protein-protein interaction false discovery rate (PPI-FDR) was imposed on each of the datasets and together 560 protein interactions are reported at a combined FDR of 2.5% (Lenz *et al*, 2021) (**Supplementary Table S1**). These 560 PPIs are underpinned by 1268 unique residue pairs. The interaction network contains 337 unique proteins, with a further 629 proteins detected with only self-links. This is a substantial fraction of the 1982 proteins detected in a whole-cell analysis using standard proteomics (**Supplementary Table S2**). Protein abundance was a key factor for a protein to be detected with crosslinks, with the median abundance of crosslinked proteins being about a magnitude higher than that of all detected proteins (iBAQ 2.7 × 10^8^ compared to 1.8 × 10^7^ **Fig. S1B**).

Of the 560 protein interactions detected by crosslinking, 176 are previously reported in *Subti*Wiki, with 384 remaining as not previously identified. As has been seen in other studies, some particularly abundant proteins contribute many interactions to whole-cell crosslinking MS approaches (O‘Reilly *et al*, 2020; Chavez *et al*, 2016). The highly abundant ribosomal proteins L7/L12 (RplL), L1 (RplA) and RS3 (RpsC), the elongation factors Ef-Tu (TufA) and Ef-G (FusA), and the RNA chaperones CspC and CspB, are identified crosslinking to more than 20 proteins each. Each of these proteins, aside from CspB, are in the top 30 proteins by intensity, with Ef-Tu (TufA) and RplL being the two most intense (**Supplementary Table S2**). If the interactions with these proteins are removed, this leaves 310 interactions, among them 186 novel interactions (**Figure 1A and S1D**). Checking the consistency of our data with known structures, we mapped the crosslinks in the dataset on the known structure of the *B. subtilis* RNA polymerase and homology models of the DNA gyrase and ATP synthase. 95 out of 98 crosslinks on these complexes were within the expected 30 Å distance between the Cα atoms (**Fig. S2**). Crosslinks mapped onto the ribosome showed 74 of 343 (21.5%) crosslinks were overlength, but these could come from multiple different states of ribosomes present in the cell, including multi-ribosome interactions and pre-ribosome assemblies (**Fig. S2**).

**Figure 1.**
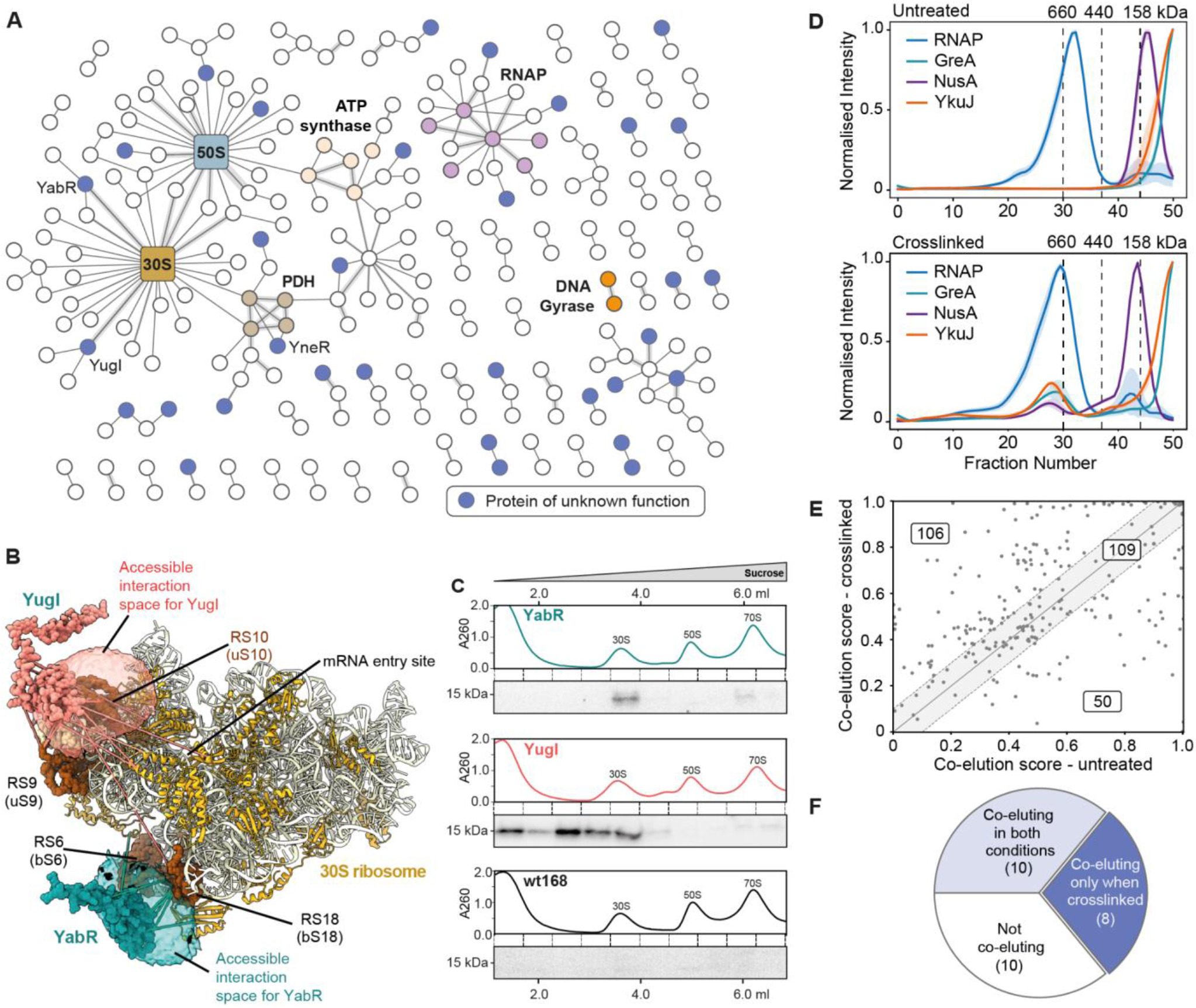
Whole-cell crosslinking reveals protein-protein interactions otherwise lost upon cell lysis. **A-** PPIs identified at 2% PPI-level FDR (interactions to seven abundant and highly crosslinked proteins are removed for clarity). Previously uncharacterized proteins are shown in blue. Selected complexes are highlighted. **B-** The accessible interaction space of YugI and YabR to the 30S ribosome calculated by DisVis (van Zundert & Bonvin, 2015). The volumes represent the positions consistent with 10 of 14 detected crosslinks for YugI and 6 of 8 crosslinks for YabR, indicating the location of their binding sites on the 30S ribosome. **C-** Sucrose gradient (10-40% w/v) of *B subtilis* lysate separating the 70S, 50S and 30S ribosomes from smaller proteins and their complexes. Western blots show that his-tagged YabR and YugI (both ∼15 kDa) co-migrate in the sucrose gradient with the 30S ribosome, the control, wild type *B. subtilis* 168, does not. **D-** Smoothed elution profiles from the CoFrac-MS analysis of the RNA polymerase (RNAP) and the known binders GreA and NusA and the uncharacterized protein YkuJ. The mass spec intensity is normalized maximum across fractions, and averaged across replicates and subunits. Top: untreated cells; Bottom: crosslinked cells. One standard deviation from the mean per fraction is shaded. The interaction with YkuJ is stabilized upon crosslinking prior to fractionation. **E-** PPIs detected by crosslinking MS analyzed by CoFrac-MS. Cofractionation measured by PCprophet co-elution scores in the crosslinked (y-axis) and untreated (x-axis) condition. PPIs within the dashed lines were considered equally predicted in both conditions (data in Table S4). **F-** Annotation of cofractionation behavior of uncharacterized proteins and their binding partners for protein pairs identified by crosslinking MS. Ribosomal proteins and proteins with missing CoFrac-MS data in either the crosslinked or the untreated CoFrac-MS condition were removed.

From the many proteins that were found with crosslinks to ribosomal subunits, two interactors stood out with crosslinks to multiple 30S proteins in close proximity; YugI with a total of 34 links to eight 30S proteins and YabR with 10 links to four proteins (**Fig. S3**). YugI has been previously detected as a binder of the 30S subunit (Natori *et al*, 2007), albeit its putative binding site remains unknown. YabR is uncharacterized. They are conserved paralogs in Firmicutes, containing an S1 RNA binding domain and share 51% sequence identity, but crosslink to different surfaces of the 30S ribosomal subunit (**Fig. 1B**). We pursued these interactions by constructing a strain that expresses C-terminally His-tagged YabR and YugI at their native loci in *B. subtilis*. Both YugI and YabR co-migrate specifically with the 30S subunit of the ribosome in a sucrose gradient (**Fig. 1C, Fig. S3**). Bacterial-two-hybrid assays were performed to test the interaction of YabR and YugI with the ribosomal proteins to which the most crosslinks were detected, namely S6 (bS6/RpsF) and S18 (bS18/RpsR) for YabR, and S2 (uS2/RpsB) and S10 (uS10/RpsJ) for YugI (**Fig. S3**) (Ban *et al*, 2014). The assay confirmed an interaction of YabR with S18 and YugI with S10. Furthermore, a *yugI* deletion strain showed increased resistance to the translation inhibitor tetracycline, supporting a functional link of the protein and the ribosome (**Fig. S3**).

Together with YabR we observed a total of 33 uncharacterized proteins in our crosslinking MS network. We wondered why we could identify these in PPIs which were missed in previous studies. To probe if the fixing of these interactions in the cell by crosslinking was key, we next investigated which protein complexes survived cell lysis and chromatographic separation by size exclusion chromatography (SEC) from untreated and from crosslinked cells.

### Crosslinking stabilizes interactions for identification by Cofractionation MS

In a second approach to detect PPIs, the soluble proteomes of both crosslinked and untreated *B. subtilis* cells were fractionated by SEC. 50 fractions were collected and analyzed by quantitative LC-MS (**Supplementary Table S3**). The subunits of several known complexes co-eluted nicely, for example the RNA polymerase, 50S ribosome and the stressosome (Kwon *et al*, 2019) **(Fig. S4**). Crosslinking stabilized some members of complexes and aided their co-elution. For example, the known RNAP binders NusA and GreA were only found eluting with the RNAP when stabilized by crosslinking (**Fig. 1D**), whereas the subunits of the core RNAP co-elute in both conditions (**Fig. S4**).

Interestingly, the uncharacterized protein YkuJ was observed co-eluting with these RNAP binders exclusively when first stabilized via crosslinking in cells. To evaluate the general benefit of crosslinking in stabilizing protein complexes prior to cell lysis, we analyzed the co-elution behavior of PPIs that were identified by crosslinking MS. Their co-elution scores were computed using PCprophet (Fossati *et al*, 2021), as described in the Methods. After filtering out intra-ribosome pairs, 109 of 265 PPIs displayed similar co-elution behavior if crosslinked or not. Of the remaining 156 PPIs, more than two thirds had a higher co-elution score following in-cell crosslinking (**Fig 1E**).

Stabilizing the proteome prior to cell lysis is especially powerful for identifying interactions involving uncharacterized proteins. We manually inspected the co-elution behavior of PPIs identified by crosslinking MS involving at least one uncharacterized protein in our CoFrac-MS data (**Fig S5**). Of these 39 PPIs, 28 could be compared, as there was data available for both crosslinked and untreated conditions. Ten PPIs showed no co-elution or were hard to classify, ten co-eluted in both conditions, and eight co-eluted or showed dramatically better co-elution only in the crosslinked condition (**Fig 1F**). Thus, crosslinking increased the percentage of identified PPIs involving uncharacterized proteins in the CoFrac-MS data from 36% to 64%.

For the generation of PPI candidates via CoFrac-MS we prepared the crosslinked and the untreated dataset by filtering them to proteins with high abundance in each of the three replicas (see methods) and removing ribosomal proteins, as lysis conditions were not selected for ribosome stability. We then analyzed co-elution behavior in PCprophet and thresholded our data to a co-elution score cutoff of 0.8 to retain only the highest confidence candidate interactions, as shown in the receiver operator characteristic curve **(Fig. S6)**. Due to the limited resolution of the column, many proteins were calculated as co-eluting in the final fractions (molecular weight <200 kDa) and near the void volume. These were removed from our data by excluding groups with more than ten members. The members of the remaining groups of co-eluting proteins were permuted all-against-all within each group into binary interactions for further analysis. This basic co-elution analysis resulted in 667 candidate PPIs total, with 449 from crosslinked cells and 318 from the untreated cells (**Fig. 1D, Fig. S6A, Supplementary Table S4**). Some proteins were only detectable from the untreated cells - this may be due to the crosslinking making them insoluble or linking them together into particles that were too large to be separated on this SEC column. The candidate PPIs from CoFrac-MS were very complementary to the crosslinking MS data, increasing the total number of our PPIs and candidate PPIs to 878, with only 4% overlap between the two techniques. The newly discovered PPIs have been added to the *Subti*Wiki database (Pedreira *et al*, 2022) (see methods).

### A system-wide PPI candidate list

To generate a comprehensive PPI candidate list for system-wide structure modeling with AlphaFold-Multimer, we added known PPIs that lack structural information to our experimentally identified PPIs. We downloaded the high-confidence protein interactions from the *Subti*Wiki database (2615 total), which are derived from various techniques, including two-hybrid screens and co-purification (Marchadier *et al*, 2011; Meyer *et al*, 2011; Commichau *et al*, 2009). The *Subti*Wiki database is manually curated, and should be enriched for direct interactions. From this list, we removed the intra-ribosome interactions due to the large amount of rRNA that complicates PPI structure prediction. We further removed homodimers and those having homologs in the Protein Data Bank (PDB) (sequence identity > 30% and BLAST EValue < 10^−3^), yielding a final list of 1218 previously known PPIs with no high-quality structural information. Similarly, we also filtered candidate PPIs from our experimental approaches to remove intra-ribosome interactions. This resulted in a final combined list of 2032 candidate PPIs for submitting to AlphaFold-Multimer (**Fig. 2A**). Surprisingly, the overlap between the three datasets is limited (**Fig. S6C**), testifying to the complementarity of approaches.

**Figure 2.**
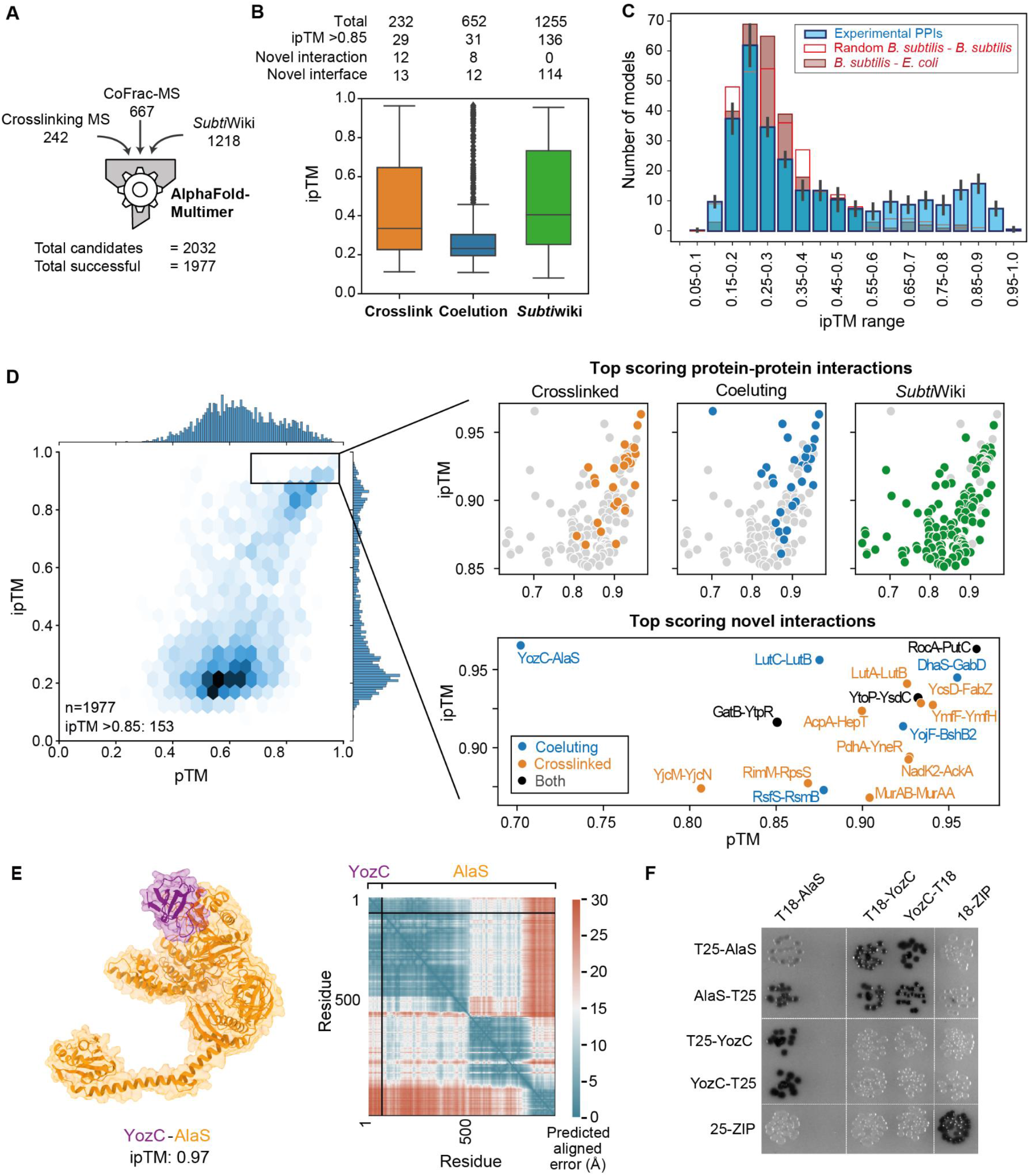
Structure prediction of binary complexes with AlphaFold-Multimer. **A-** The 1977 predicted PPIs for AlphaFold-Multimer interface prediction from crosslinking MS, CoFrac-MS and *Subti*Wiki. **B-** Breakdown of AlphaFold ipTM score distributions by PPI origin. Annotation of score distributions for PPIs annotated by being present in the PDB (seq. identity > 30% and Evalue < 10^−3^) or by their presence in STRING (combined score > 0.4). “Novel interaction” refers to a previously unknown PPI, while “novel interface” refers to the lack of homologous structures for the PPI in the PDB. **C-** Noise model evaluation of ipTM distribution of AlphaFold PPIs. Subsamples of 300 PPIs from our datasets (target distribution) are compared to 300 PPIs made up of random *B. subtilis* proteins from the PPI candidate list combined with random proteins from the *E. coli* genome (noise distribution). While targets show a bimodal distribution, indicating the high confidence of models with ipTM > 0.85, the noise distribution is one-tailed, approximating the likelihood of random interface prediction in the various ipTM ranges. **D-** The 1977 protein-protein interactions (PPIs) modeled by AlphaFold-Multimer distribute over the full pTM and ipTM range, with a subpopulation of highly confident predictions with ipTM > 0.85. Insets showing high-ranking models colored by dataset of origin, and the top-ranking PPIs not previously annotated in *Subti*Wiki **E-** A novel PPI from the co-elution dataset showing the alanine tRNA synthetase subunit AlaS interacting with the uncharacterized protein YozC. The high ipTM value is reflected in the predicted aligned error plot, which also shows that the C-terminal region of AlaS, not involved in the interaction, is flexible with respect to the YozC-AlaS module. **F-** Bacterial-2-hybrid assay to validate the interaction between YozC and AlaS. N- or C-terminal fusions of YozC and AlaS to the T18 and T25 domains of the adenylate cyclase CyaA were created and tested for interaction in the *E. coli* strain BTH101. Colonies turn dark as a result of protein interaction, which leads to the restoration of the adenylate cyclase activity and therefore expression of the ß-galactosidase. A leucine zipper domain was used as a positive control.

### Identification of protein-protein interaction interfaces by AlphaFold-Multimer

We derived structural models of these PPIs by submitting each protein pair to AlphaFold-Multimer (version 2.1), which uses a model trained on the protein structure database and multiple sequence alignments to infer the structure of proteins and multiprotein complexes (**Supplementary Table S5**). The resulting 1977 models were assessed for overall predicted TM-score (pTM), and interface predicted TM-score (ipTM) (**Fig. 2B,D**). These two error metrics rely on estimating the overall similarity of the model to the unknown true solution by predicting the TM-score (Zhang & Skolnick, 2007) of the two structures on all residues (pTM) or on inter-subunit distances only (ipTM). Thus, pTM reports on the accuracy of prediction within each protein chain, and ipTM on the accuracy of the complex. A TM-score of 0.5 is broadly indicative of a correct fold/domain prediction (Zhang & Skolnick, 2007; Xu & Zhang, 2010; Andreeva *et al*, 2020; Sillitoe *et al*, 2021), while scores above 0.8 correspond to models with matching topology and backbone path (Kufareva & Abagyan, 2012; Olechnoviĉ *et al*, 2019; Xu & Zhang, 2010). An ipTM > 0.85 has proven reliable in other analyses when compared to known interface TM-score and the DockQ docking quality score (Evans *et al*, 2022; Burke *et al*, 2021; Bryant *et al*, 2022b). In total, the predictions resulted in 153 high-confidence PPI models (ipTM > 0.85) (**Fig. 2D**). This includes 17 novel interactions for which no annotation had been previously available (**Fig. 2D, S7**), and 130 interactions with no good template homologous structures in the PDB i.e. for which we predict a first high-quality model even though many have previously been annotated in *Subti*Wiki and thus worked on. A further 396 models have a lower confidence (ipTM 0.55-0.85), 26 of which represent novel interactions. The candidates from the three approaches yielded different subsets of high-scoring models, with crosslinking MS providing the highest ‘hit rate’ for structural modeling of novel PPIs (12% of crosslinking MS PPIs lead to models with ipTM >0.85, 4% of CoFrac-MS, and 11% of the *Subti*Wiki dataset). (**Fig. 2B**). This agrees with co-elution not selecting for direct binary interactions and thus giving the lowest hit rate. In contrast, manual curation of the available literature and crosslinking MS yield comparable outcomes.

We set a stringent cutoff of ipTM=0.85 for calling high-confidence PPI models. In order to prove the robustness of our score cutoff, we employed a noise model in which 300 *B. subtilis* proteins from our datasets were predicted as pairs with random *E. coli* proteins. The ipTM distribution of the resulting decoy PPIs was compared with 10 subsamples of our AlphaFold-Multimer predictions (**Fig. 2C**), showing that ipTM < 0.55 for AlphaFold-Multimer indicates a random prediction, while 0.55-0.85 performs better than random, with increasing accuracy. No decoy PPIs reported an ipTM > 0.85. We take this result to indicate that, especially in the ipTM range 0.55-0.85, AlphaFold-Multimer models require additional validation by other experimental approaches.

Each predicted protein-protein interaction was also assessed in terms of its predicted aligned error (PAE) matrix, which reports on the predicted error in the position of a residue if the protein were aligned to the true solution elsewhere along the sequence. PAE can be used to estimate confidence in positions of parts of the protein or complex relative to the rest. In the example shown in **Fig. 2E**, the novel interaction identified by CoFrac-MS between the alanine-tRNA synthetase AlaS and the uncharacterized protein YozC is shown. The model has the highest ipTM score (0.97) in the dataset, but a low pTM score (0.70), indicating high confidence in the interface but a lower confidence in the prediction of the overall structure. The PAE plot shows that the relative position of YozC and the AlaS N-terminal region has a very low predicted aligned error, but the position of these two regions relative to the rest of AlaS, which contains two more domains, is uncertain. To confirm this interaction, we performed a bacterial two-hybrid experiment that demonstrated that these proteins directly interact (**Fig. 2F**).

It is important to note that predictions with low ipTM values indicate poor models, but do not necessarily mean the two proteins do not interact. AlphaFold-Multimer can provide inaccurate results in cases where the protein pair resides in a larger complex, where the interaction is mediated by nucleic acids or other molecules, as well as in cases where the interaction is dependent on a post-translational modification.

### Validation of AlphaFold-Multimer models by crosslinking MS

Our high-quality crosslink data provides insights into the structure of protein complexes inside cells and allows validating the corresponding AlphaFold model (**Fig. 3A**). We found a strong correlation between ipTM and restraint satisfaction of heteromeric crosslinks, despite the fact that crosslinking information was not used in AlphaFold model prediction. Crosslink violation is especially low with ipTM 0.85, indicating that high-confidence models agree with the residue-residue distances observed *in situ*.

**Figure 3.**
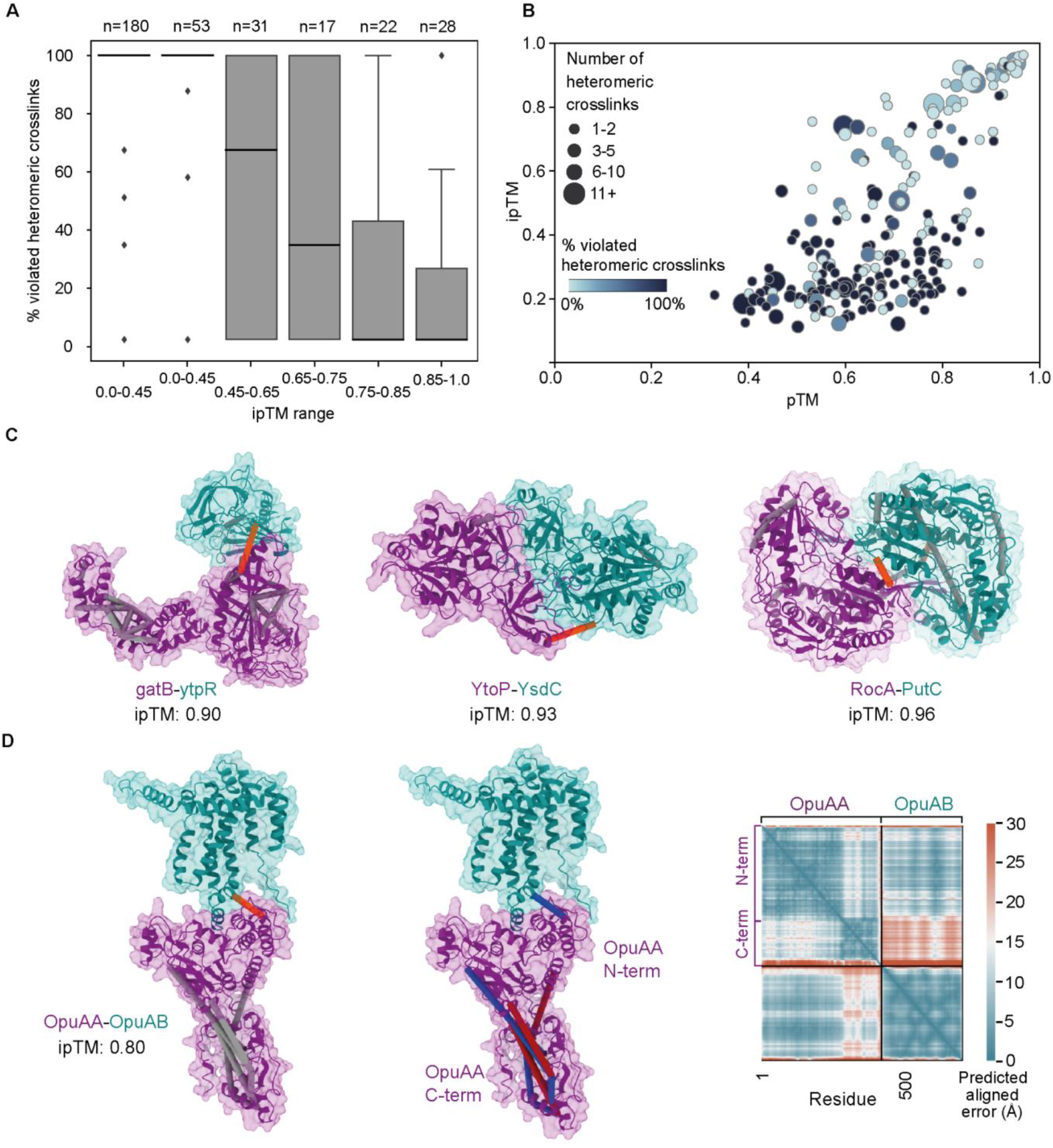
Crosslinking MS validation of AlphaFold-Multimer models. **A-** Percentage of heteromeric crosslink restraint violation per range of ipTM. **B-** Bubble plot showing numbers of heteromeric crosslinks violated for each PPI identified by crosslinking MS against the ipTM and pTM distribution. **C-** Successful predictions consistent with crosslinking MS, including predictions of paralogs (YtoP-YsdC, RocA-PutC). Self crosslinks in gray, heteromeric crosslinks in orange. **D**-Crosslinks highlighting flexibility within the OpuAA-OpuAB dimer. The OpuAA N-terminal domain is predicted with a high pAE to the C-terminal region. The crosslinks corresponding to these interdomain distances are also violated, indicating flexibility between these 2 domains. Left: Self crosslinks in grey, heteromeric crosslinks in orange; center: satisfied crosslinks (<30 Å Cα-Cα) in blue, violated crosslinks in red; right: predicted aligned error plot.

In the ipTM range of 0.55-0.85 models show a wide distribution of heteromeric restraint violation percentages (**Fig. 3A**), indicating that models in this ipTM range may be independently validated or at least partially rejected based on experimental information. A low degree of restraint violation suggests that the conformations predicted are at least in some features representative of the structures inside cells. High restraint violation may indicate the model does not reflect the in-cell conformation in the regions covered by crosslinking MS data, or that the prediction is far from the true solution. Nevertheless, crosslinking MS data show that models in the ipTM 0.75-0.85 range are more likely to be consistent with *in situ* structural restraints than models in the 0.55-75 range, indicating increasing model quality (**Fig. 3A**). It is also noteworthy that the models with low (<0.55) ipTM display a median 100% violation rate of heteromeric crosslinks (**Fig. 3A**), corroborating the poor nature of interfaces in models with low ipTM scores.

Match to crosslinking MS data can therefore independently confirm predicted interfaces, especially for those PPIs with a high number of heteromeric crosslinks (**Fig. 3B**), where a large swath of the interface is covered by crosslinking MS data. For example, the crosslinking MS data confirms the predicted model for the novel interaction between the B subunit of the glutamyl-tRNA amidotransferase (GatB) and the uncharacterized protein YtpR, which has putative RNA-binding activity (**Fig. 3C**). Several crosslinks within GatB additionally validate the topology of this protein’s fold.

Self crosslinks may also provide important insights into protein conformation, as they may be used to indicate which models below our ipTM threshold are reliable, as in the case for the membrane transporter subunits OpuAA-OpuAB (ipTM=0.80). Here, heteromeric crosslinks validate the predicted interface and self crosslinks highlight the flexibility of the OpuAA N-terminal region with respect to the rest of the complex, which can be also seen in the predicted aligned error plot (**Fig. 3D**).

Due to its sequence resolution, crosslinking MS can also provide information on the interaction of paralogs for which so far only homomeric complexes have been reported. In our high-scoring models we had four dimers of paralogs; RocA-PutC, YtoP-YsdC, YmfF-YmfH, and MurAA-MurAB. In the case of RocA-PutC (**Fig. 3C**), both proteins are paralogs of *Bacillus halodurans* 1-pyrroline-5-carboxylate dehydrogenase, which has been solved as a homodimer (PDB: 3qan), with sequence identities of 69% and 74% respectively. Due to the high sequence identity, AlphaFold templates RocA-PutC on the homomeric *B. halodurans* RocA1-RocA1 complex (PDB: 3qan), leaving unclear if the heteromeric model is physiologically relevant. Multiple residue-residue pairs are detected for RocA-PutC, clearly indicating the heteromeric complex is formed *in situ*. The crosslinks are satisfied in the AlphaFold model, confirming the interface. Moreover, no crosslinks indicating a homodimer (involving the same peptide pair) were observed.

### Inferring novel protein complexes from binary prediction

Due to the intrinsic limitations of any network analysis, all three approaches used to generate binary PPIs here (crosslinking MS, CoFrac-MS and the *Subti*Wiki database) can provide only indirect information on higher-order interactions. The binary interactions predicted above can be independent binary events or be part of larger multiprotein complexes. Such assemblies may contain many copies of the two proteins or involve additional subunits. Nevertheless, the binary interactions can be used to infer associations in larger assemblies.

To look for potential higher order complexes in our binary PPI structure predictions, we plotted all PPI predictions with ipTM > 0.65 as a network (based on **Fig. 2C**). Groups of predicted PPIs might indicate higher order complexes. In total 64 groups were identified. These ranged from those containing only three proteins, to the largest containing 16 members (**Fig. S8**). It is of interest to note that the two largest potential complexes each contain functionally related proteins that are involved in DNA replication and recombination (centered around DnaN) (Lenhart *et al*, 2012) and in sugar transport by the phosphotransferase system (PTS, centered around PtsH) (Stülke & Hillen, 1998). In the case of the PTS interactions, most of them are known binary interactions involved in phosphotransfer of one protein to the other or in binary regulatory interactions. Thus, a large complex is not likely for the PTS proteins, whereas the formation of one or two large complexes is feasible for the replication and recombination proteins. For large clusters of interacting proteins there are many potential combinations of stoichiometries that could be predicted, and so prior knowledge is required to model complexes correctly (Gao *et al*, 2022; Bryant *et al*, 2022a). In order to simplify the problem for the purpose of this study, we predicted only potential heteromeric trimers with a 1:1:1 stoichiometry. Our network identified 33 groups of only three proteins, including 5 potential complexes involving novel interactions (**Fig. 4A**).

**Figure 4.**
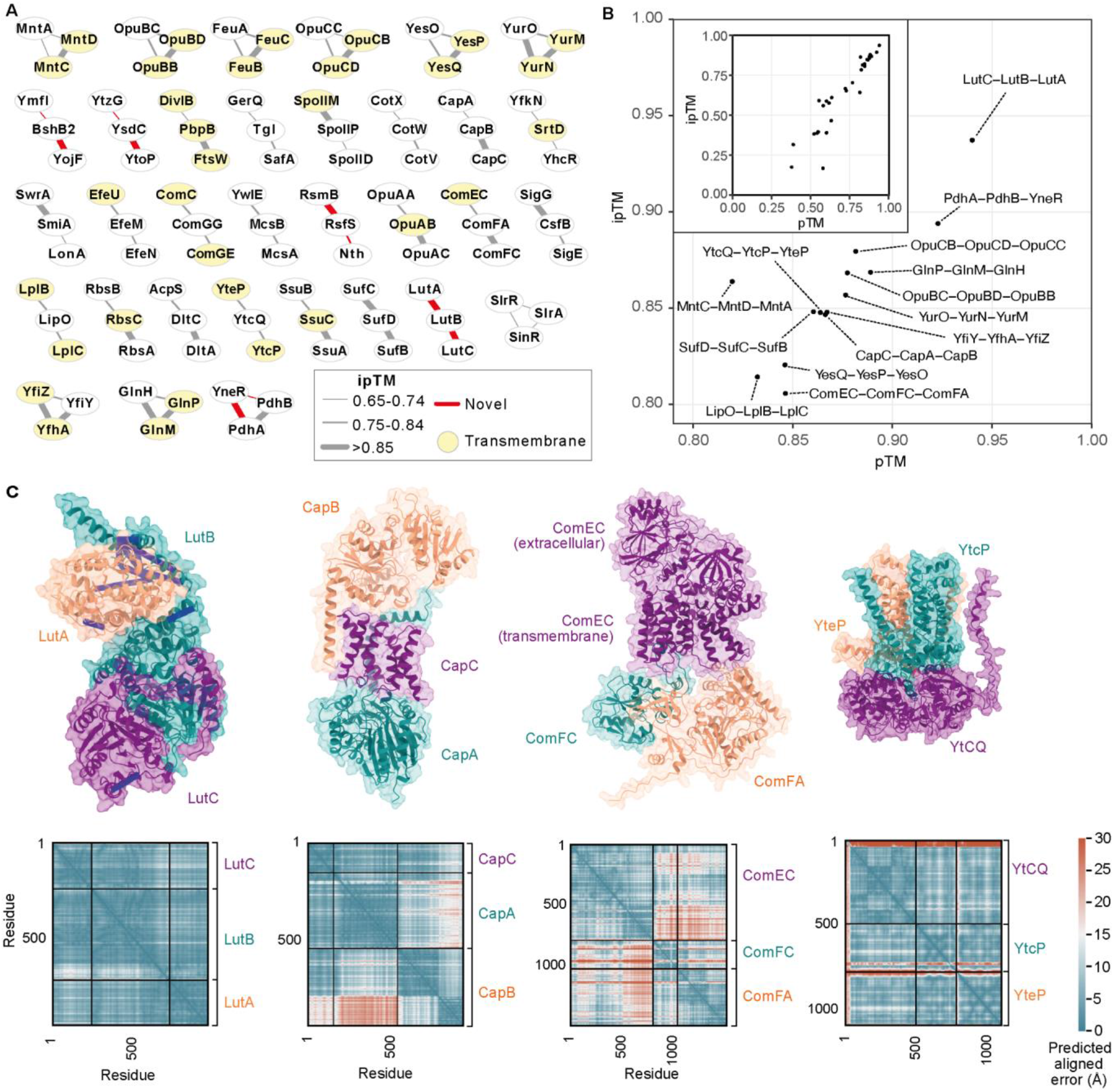
Building complexes from binary interaction predictions. **A**. All dimeric PPIs with predicted ipTM >0.65 which form connected groups of only 3 proteins are shown. **B**. The 33 candidate 1:1:1 trimers PPIs modeled by AlphaFold-Multimer (version 2.2.1) distribute over the full pTM and ipTM range (inset). Trimers with an ipTM > 0.80 are labeled. **C**. Selected predicted structures of trimeric complexes with ipTM > 0.80 and their associated PAE plots. Crosslinks are visualized on LutA-LutB-LutC; and satisfied crosslinks (<30 Å Cα-Cα) in blue, violated crosslinks in red.

The 33 candidate trimers were predicted with AlphaFold-Multimer (version 2.2.1) (**Supplementary Table S6**), resulting in 14 trimer predictions with ipTM > 0.8. The top-ranking hit is a previously unknown complex between the proteins of the lactate utilization operon LutA-LutB-LutC (Chai *et al*, 2009). The interactions are identified by a combination of crosslinking MS (LutA-LutB) and CoFrac-MS (LutB-LutC). In the predicted structure, the PAE plot shows a highly confident placement of the whole sequence of the subunits. LutB contains an Fe-S cluster that is located away from subunit interfaces, though the LutC N-terminal region forms extensive interactions with the LutB α2 helix covering the Fe-S site.

One of the predicted complexes is the complex between CapA, CapB, and CapC. These proteins catalyze the synthesis and the export of γ-polyglutamate (PGA), an extracellular polymer. In *B. subtili*s, all of the enzymes needed for γ-PGA synthesis are encoded in the *capBCAE* operon (Urushibata *et al*, 2002). CapB and CapC form the γ-PGA synthase complex, whereas CapA and CapE co-operate in export (Candela *et al*, 2005). The formation of a CapBCA complex has been suggested previously, with a tight interaction between the ligase subunits CapB and CapC and a loose interaction of the ligase to CapA (Ashiuchi *et al*, 2001). Our work provides evidence for the existence of the CapBCA complex with this confident structural prediction (**Fig. 4C**). Interestingly, in *B. anthracis*, the causative agent of anthrax, the *cap* operon is present on the virulence plasmid pXO2. This bacterium uses the γ-PGA capsule to protect itself from the host’s immune surveillance, which therefore is an important virulence factor (Jang *et al*, 2011; Mock & Fouet, 2001).

Among the top-ranking hits, we also find the competence proteins (ComEC-ComFC-ComFA) arranged in a membrane-spanning complex (**Fig. 4C**). The ComEC membrane nuclease binds ComFA, an ATPase involved in DNA import, and the late competence factor ComFC. These three proteins all localize to the cell poles and share a similar expression pattern across growth conditions (Kaufenstein *et al*, 2011; Pedreira *et al*, 2022). The interactions of ComFA with ComFC and ComEC have already been reported (Kramer *et al*, 2007; Diallo *et al*, 2017). A ternary complex between these proteins suggests that the energy provided by ComFA-mediated ATP hydrolysis fuels ComEC mediated uptake of single-stranded DNA molecules (Silale *et al*, 2021).

Finally, there are 10 transmembrane transporters and permeases predicted. All proteins are already known to belong to various classes of ATP-binding cassette (ABC) transporters or are annotated as putative ABC transporters. One example is the permease YtcP-YtcQ-YteP (**Fig. 4C**), a permease for complex carbohydrates (Ferreira *et al*, 2017; Ochiai *et al*, 2007). Other ABC transporters, like YclN-O-P-Q, fall into higher order assemblies (**Fig. S8**). For the latter complex, the known stoichiometry can even be gleaned from the binary predictions and the full complex can be modeled (**Fig. S9**). While these predictions are confident, stoichiometry information remains crucial in protein complex prediction.

### PdhI/YneR is an inhibitor of the E1 module of pyruvate dehydrogenase

The interaction of the uncharacterized protein YneR, here renamed PdhI, with the E1 module of pyruvate dehydrogenase (PdhA-PdhB) was identified by crosslinking MS. The predicted ternary complex shows a confident arrangement of the 3 proteins (ipTM=0.89), despite a low-confidence prediction of the binary PdhI-PdhB interaction. The 10 predictions could be grouped into two distinct possible configurations of the PdhA-PdhB subcomplex, which are consistent with the known ‘dimer of dimers’ stoichiometry of the E1 module (**Fig. 5A-C**). The crosslinks to PdhI were only satisfied in the worse scoring trimer conformation (**Fig. 5D**). Indeed, both high- and low-scoring predictions map to arrangements occurring in the homologous structures. Once taking both dimers into account, it is possible to use the AlphaFold models to reconstruct the full E1 PDH bound to PdhI (**Fig. 5E**). CoFrac-MS data shows PdhI co-eluting with large assemblies comprising both PdhA and PdhB, further confirming the interaction of this protein with the assembled E1 PDH module (**Fig. 5G**).

**Figure 5.**
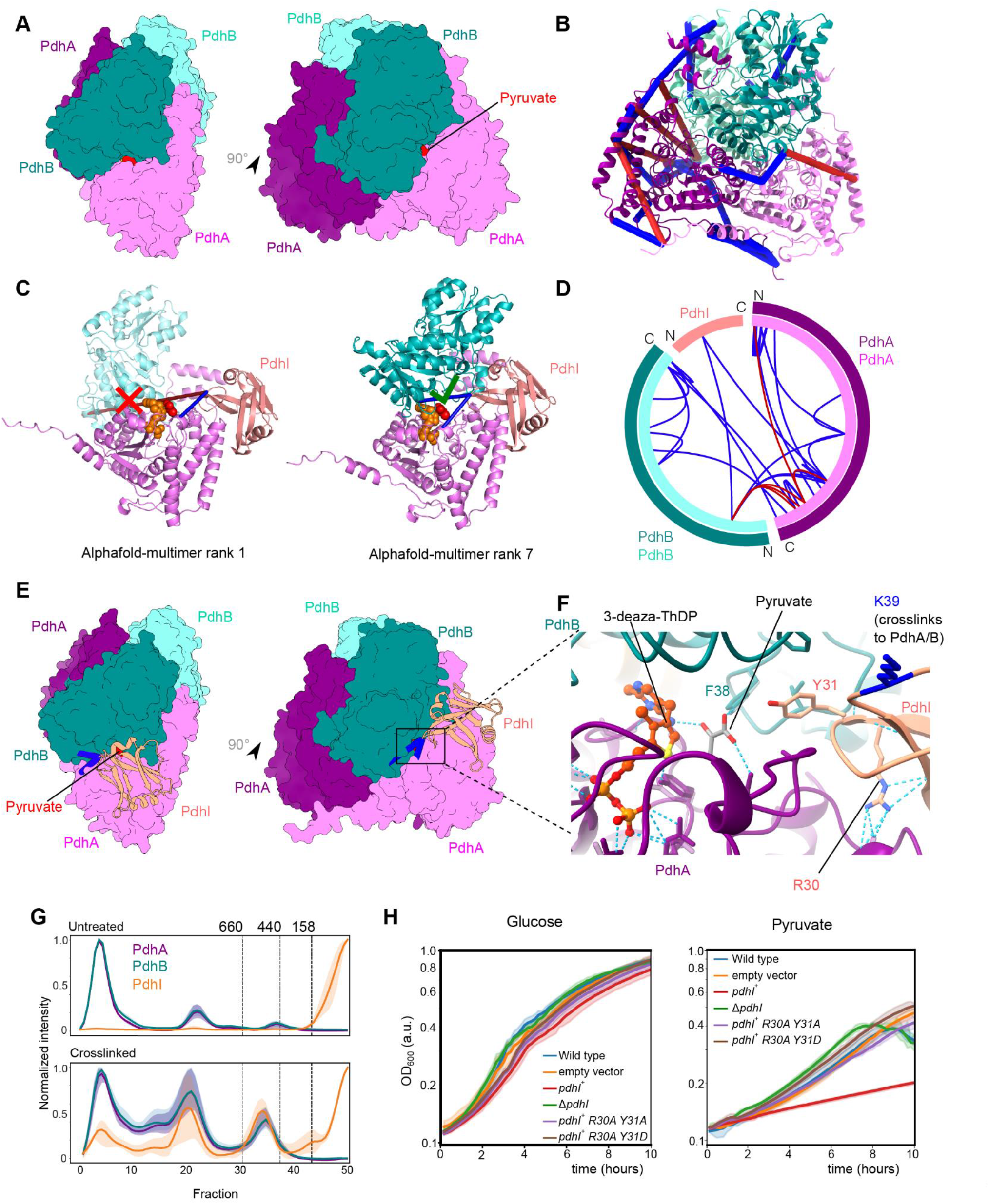
PdhI/YneR is an inhibitor of the E1 subunit of the pyruvate dehydrogenase. **A**- Homology model of *B. subtilis* E1 pyruvate dehydrogenase (PDH) based on the *Geobacillus stearothermophilus* E1p structure (PDB id 3dv0) (Pei *et al*, 2008) in surface representation. The space-fill model of pyruvate is located in the active site based on the template structure. The E1 PDH is a dimer of dimers of the PdhA and PdhB subunits, with the active site formed at the interface between a PdhA and a PdhB copy. **B**- Mapping of crosslinks onto the E1 PDH model derived from combining AlphaFold-Multimer models. Satisfied crosslinks (<30 Å Cα-Cα) in blue, violated crosslinks in red. **C**- AlphaFold-Multimer predictions for PdhA-PdhB-PdhI/YneR. The top-ranked solution by ipTM (0.89) describes the PdhA-PdhB subcomplex that does not make up the active site, while the 9th-ranked solution (0.81) identifies the active site interface. Crosslinking data clarifies the interactions between PdhI and PdhA/B. Pyruvate and 3-deaza-TdHP shown as space-fill models. Crosslink coloring as in B. **D**- Circle view of crosslinking MS data mapped onto the E1 PDH-PdhI/YneR model derived by combining AlphaFold solutions onto the known stoichiometry. Satisfied crosslinks (<30 Å Cα-Cα) in blue, violated crosslinks in red. **E**- PDH-PdhI model constructed from AlphaFold-Multimer models of the PdhA-PdhB-PdhI trimer. PdhI/YneR binds at the pocket opening onto the active site. **F-** Visualization of the active site in the AlphaFold-Multimer model (solid cartoon) with ligand positions derived PDB id 3dv0 (transparent cartoon and sticks). PdhI/YneR occludes the entrance to the active site by inserting Y31 into the pocket used for entrance of the lipoate cofactor that comes to reduce the thiamine ring in the enamine-ThDP intermediate. The original structure was solved in the presence of the enamine-ThDP analogue 3-deaza-TdHP (Pei *et al*, 2008). Key residues for ligand coordination are predicted in the same conformation by AlphaFold-Multimer. **G**- CoFrac-MS data showing co-elution of PdhA, PdhB and PdhI. The shaded area corresponds to the standard deviation between replicas. **H**- Growth curves on glucose and pyruvate. Growth experiment of wild type (blue) *B. subtilis*, PdhI/YneR overexpression (green) and PdhI/YneR knockout Δ*yneR* (red) in MSSM minimal medium with 5 mM KCl comparing growth on either glucose or pyruvate as a sole carbon source. Empty vector control in orange. Mutations in residues involved in PdhI binding to PdhA/PdhB lead to phenotypic recovery. Lines represent the mean. The shaded area corresponds to 95% confidence intervals.

In the complex, PdhI partially occludes the active site of the PdhA-PdhB dimer (**Fig. 5F, Fig. S10A**). AlphaFold predicts that Y31 of PdhI (pLDDT 79.5) inserts in the active site along the hydrophobic cavity surrounding the active site, covering the entrance to the active site. However, the prediction of this region of the complex indicates some degree of uncertainty or flexibility, as reported by pLDDT scores range from 65 to 80 in the loops forming contacts between PdhA and PdhI (**Fig. S10B**). PdhA residues 273-287, which form an extended loop in proximity of PdhI, are not resolved in the *Geobacillus stearothermophilus* E1p structure (PDB: 3dv0) (Pei *et al*, 2008), corroborating the flexibility of this region. Due to symmetry, it is possible that PdhI may also bind the E1 subunit in a 2:2:2 complex, though PdhI is far less abundant than E1 and is therefore likely to bind substoichiometrically (**Table S2, Fig. 5G**).

This configuration suggests that PdhI would modulate the activity of the E1 subunit. To test this, we generated two strains, one that overexpressed PdhI and one with PdhI knocked out (**Fig. 5H**). These strains did not have growth defects compared to the WT when grown with glucose as the main carbon source. However, cells with overexpressed PdhI had a dramatic growth defect when grown with pyruvate as the sole carbon source, indicating that PdhI acts as an inhibitor of pyruvate dehydrogenase. Strains overexpressing PdhI carrying mutations at Y31 and R30 (which forms hydrogen bonds to PdhA) do not show growth defects on pyruvate media, confirming the interface predicted by AlphaFold is critical to the function of PdhI (**Fig. 5H**).

## Discussion

*B. subtilis* is a model Gram-positive bacterium, with extensive genetic data (Michalik *et al*, 2021) and its protein structures modeled to a high degree of accuracy (Varadi *et al*, 2022). Nevertheless, 25% of proteins in *B. subtilis* remain poorly characterized or even lack any characterization (Michna *et al*, 2015). In this paper, we describe genetic-free approaches for protein-protein interaction screening capable of producing large numbers of novel protein-protein interactions along with their topologies by fixing interactions in cells. The experimental approaches yielded 44 high-quality PPI models (ipTM 0.85). Adding interactions curated in *Subti*Wiki led to high-quality models for 114 binary interactions with no previous good structural homology. Considering only 601 non-ribosomal *B. subtilis* PPIs had previous structural information, mostly from homology, this is a substantial increase of the structural coverage of the known interaction space. Our approach is particularly successful for membrane proteins, which represent a challenge for structural and systems biology methods. 80 of our 153 high-quality dimers include proteins with transmembrane domains, and membrane proteins are present in half of our predicted trimer structures.

In addition to highly confident models (ipTM > 0.85), the AlphaFold PPI models in this study can be classified into those that cannot be confidently predicted as a protein pair (ipTM < 0.55), and the “gray zone” of models with an intermediate ipTM range, based on the noise model for error rate determination employed in **Fig. 2C**. These boundaries are due to change as deep learning prediction develops, and we believe modeling the chance of random predictions will be beneficial also in future PPI screens. High-scoring models display very high crosslink distance restraint satisfaction, showing the accuracy of high-ipTM predictions (**Fig. 3**). For models of intermediate confidence, ipTM alone cannot distinguish reliably between trustworthy and random. However, experimental structural data such as those offered by crosslinking MS may provide crucial evidence and offer a systematic path to expanding the reliability of AlphaFold into lower ipTM scores.

It is important to note that models with low ipTM do not necessarily mean that these are not true interactors. This is exemplified in our data, where the novel ribosome binding proteins YabR and YugI had their best predictions to RS11 and RS2, with ipTM of only 0.53 and 0.33, respectively (**Supplementary Table S5**). These proteins had novel interactions to multiple 30S ribosome proteins detected by crosslinking MS. This interaction may be mediated by the rRNA elements located in the proximity of the interacting partners, especially given the presence of RNA binding domains in both YugI and YabR.

In this work, we have identified several novel interactions that are likely of biological relevance. For example, the previously uncharacterized protein YtpR was found in complex with the B subunit of the glutamyl-tRNA amidotransferase (GatB). The YtpR protein contains a tRNA-binding domain at its C-terminus. It is tempting to speculate that it presents the tRNA^Gln^ preloaded with glutamate to the GatCAB complex to convert the glutamate cargo to glutamine. Interestingly, the YtpR protein is highly expressed in *B. subtilis* and is ubiquitous in archaea and bacteria which use the Gat-dependent pathway for the synthesis of tRNA^Gln^ (Nakamura *et al*, 2006). Taken together, this suggests that the interaction between YtpR and GatB is highly conserved among prokaryotic organisms and functionally relevant.

We also predicted the previously uncharacterized protein PdhI/YneR in complex with PdhA and PdhB, which make up the E1 module of the pyruvate dehydrogenase complex. The predicted binding interface near the active site, confirmed by crosslinking MS, led us to hypothesize that PdhI is a negative regulator of the E1 module. Indeed, PdhI overexpression dramatically slowed growth on pyruvate as the sole carbon source. The predicted insertion of PdhI into the hydrophobic cavity that surrounds the active site of the enzyme immediately suggests the molecular mechanism for the control of pyruvate dehydrogenase activity by PdhI. Moreover, site-directed mutagenesis confirmed the predicted molecular mechanism and the site of interaction. This example demonstrates the power of combining global proteomic approaches to identify PPIs with artificial intelligence-assisted structure prediction and experimental validation to uncover the function of so far unknown proteins.

Crosslinking MS holds the potential to capture all PPIs *in situ*, but current technology limits the depth of analysis that can be reached. Thus, we complemented it here with the noisier CoFrac-MS. These approaches are scalable, are in active development (McWhite *et al*, 2020; Rosenberger *et al*, 2020; Bludau *et al*, 2021; Chavez *et al*, 2018) and can be applied to any species or cell type. Our large-scale hybrid PPI screen followed by AlphaFold-Multimer structure prediction led to high-quality models for PPIs comprising several uncharacterized proteins, for which we provide association partners. It is possible to predict multisubunit complexes *de novo* from the binary interactions by combining pairwise predictions (Fig. S9) (Bryant *et al*, 2022a, 2022b). Principally, the binary models of AlphaFold may provide a starting point for reconstructing models of larger protein complexes. Predicting complexes using the correct stoichiometry of a complex, like in the case of the E1 PDH, can improve ipTM (Gao *et al*, 2022). Yet, when stoichiometries are unknown, the results are difficult to interpret (Evans *et al*, 2022; Burke *et al*, 2021). Systematic searching of stoichiometries in protein structure prediction is an active area of research (Bryant *et al*, 2022b), and experimental efforts to determine stoichiometries are collected systematically (Hu *et al*, 2019; Dey & Levy, 2021).

The combination of crosslinking MS and CoFrac-MS used in this study can accelerate the discovery of protein-protein interactions from in-cell and in-lysate data. These experimental techniques facilitate the untargeted investigation of PPIs and therefore make up one of the key approaches to identify the function of understudied proteins (Kustatscher *et al*, 2022). These PPIs, combined with previously annotated indirect interactions from databases such as *Subti*Wiki, can be employed by AlphaFold-Multimer to generate highly accurate structural models of known and novel interactions and complexes at scale. For *E. coli*, a bacterium of ∼4500 genes, it is estimated that there are 10,000 specific protein-protein interactions (Rajagopala *et al*, 2014). While exact numbers are difficult to estimate, the number of interactions considered here likely cover a substantial fraction of the interactome. This study shows the power of untargeted PPI mapping approaches and especially in-cell crosslinking in establishing structure-function relationships for currently uncharacterized proteins, and the potential of hybrid experimental PPI screens and structure prediction for the future of structural systems biology.

## Methods

### Materials

Unless otherwise stated, reagents were purchased in the highest quality available from Sigma (now Merck), Darmstadt, Germany. Empore 3M C18-Material for LC-MS sample cleanup was from Sigma (St. Louis, MO, USA), glycerol from Carl Roth (Karlsruhe, Germany). DSSO (disuccinimidyl sulfoxide) crosslinker from Cayman Chemical (Ann Arbor, MI, USA). Dimethylformamide (DMF) from Thermo Fisher Scientific. EDTA-free protease inhibitors (Roche) lysozyme (Sigma Aldrich), acrylamide (VWR), C18 HyperSEP cartridges (Thermo Scientific)

### Biomass production

*B. subtilis* strain 168 was grown on Luria-Bertani (LB) agar at room temperature in all steps. A single colony was transferred into LB broth and a pre-culture grown overnight. The pre-culture was diluted to a starting OD_600_ of 0.005 and grown to an OD_600_ of ∼0.6 before being harvested by centrifugation at 4500 g for 5 min. The pellets were resuspended and washed with PBS and pelleted again, twice.

‘Crosslinked cells’: cells were resuspended and crosslinked in fresh PBS at a final concentration of 5 mg wet cell mass/ml, 1.4 mM DSSO (CoFrac-MS) or 2.6 mM DSSO (crosslinking MS) and 5% DMF. Reactions were allowed to proceed for 60 min at room temperature and quenched with 100 mM ammonium bicarbonate (ABC) for 20 min. Cells were pelleted at 4°C, washed with ice-cold PBS and snap-frozen in liquid nitrogen.

‘Untreated cells’: Cells were resuspended for a third time in fresh PBS to a final concentration of 5 mg wet cell mass/ml and 5% DMF and processed identically to the crosslinked cells.

### Proteomics for protein abundance estimation

A frozen untreated cell pellet (150 mg wet cell mass) was resuspended in fresh PBS to 150 mg/ml with 0.3 mg/ml lysozyme (Sigma Aldrich) and incubated for 30 min at 37°C in a water bath. EDTA-free protease inhibitors were added just prior to lysis by sonication on ice using a Qsonica microtip probe (3.2 mm) for 30 seconds 1 second on/1 second off with amplitude 12-24%. After the first cycle 250 U/ml benzonase and 20 mM MgCl_2_ was added. After lysis the lysate was left to incubate for 30 min on ice and dithiothreitol (DTT) was added to a final concentration of 1mM.

Lysates were subsequently clarified by centrifugation for 30 min at 20,000 x g and 4°C. Protein in the supernatant was precipitated by chloroform/methanol precipitation (Wessel & Flügge, 1984). The pellet was resuspended in 6 M guanidine hydrochloride with 50 mM Tris-HCl (pH 8) before sonicating 5x for 30 s on ice with settings as before. Proteins were precipitated with the Wessel-Flügge precipitation and added to the rest of the proteome.

The precipitated proteome was resuspended in 8 M urea/100 mM ABC containing 1 mM DTT and incubated on a shaker for 15 min. The sample was spun down at 16,873 x g for 10 min and supernatant was diluted to 2 mg/ml after quantification by Bradford assay (Sigma Aldrich). The sample was reduced for 30 min by adding DTT to a concentration of 5 mM followed by an alkylation step with acrylamide at 15 mM for 30 min in the dark. The alkylation was quenched with 5 mM DTT. LysC was added in an enzyme/protein ratio of 1:200 (w/w) and incubated at room temperature for 4 hours before decreasing the urea concentration to 1.5 M using 100 mM ABC. Trypsin was added (enzyme/protein ratio of 1:50 w/w) and samples incubated for 8.5 h at 24°C before adding more trypsin (final enzyme/protein ratio of 1:25) for another 9.5 h. Digestion was quenched by acidification with trifluoroacetic acid (TFA) to pH 3.0 and peptides were cleaned up using a C18 StageTip (Rappsilber *et al*, 2007).

Eluted peptides were dried in a vacuum concentrator, resuspended in 1.6 % ACN (v/v) in 0.1% formic acid. Approximately 1 µg was injected into a Q Exactive HF Mass Spectrometer (Thermo Fisher Scientific, San Jose, USA) connected to an Ultimate 3000 UHPLC system (Dionex, Thermo Fisher Scientific, Germany). Chromatographic setup used the following LC gradient: Gradient started at 2% B to 5% B in 1 min, to 7.5% B in 2 min, then to 32.5% in 48 min, 40% B in 8 min, 50% B in 2.5 min followed by ramping to 90% B in 1.5 min and washing for 5 min. Each fraction was analyzed as a single injection over a total run time of 90 min each. The settings of the mass spectrometer were as follows: Data-dependent mode; MS1 scan at 120,000 resolution over 350 to 1,600 m/z; normalized AGC target of 250% with max. IT of 60 ms; MS2 triggered only on precursors with z = 2-7; 1.6 m/z isolation width; normalized AGC target of 90% with 40 ms max. IT; fragmentation by HCD using stepped normalized collision energies of 28, 29 and 31; MS2 scan resolution 15,000; peptide match was set as preferred and dynamic exclusion was enabled upon single observation for 30 seconds.

Mass spectrometry raw data was processed using MaxQuant 1.6.12.0 (Tyanova *et al*, 2016) under default settings with minor changes: two allowed missed cleavages; oxidation on methionine as a variable modifications; carbamidoethylation on Cys was set as fixed modification. The database used covered all 4,191 proteins listed for *B. subtilis* 168 in UniProt (Reviewed Swiss-Prot). The ‘matching between runs’ feature was disabled. Protein quantification was done using the iBAQ approach (Schwanhäusser *et al*, 2011). Raw data and search output are summarized in Table S2.

### Crosslinking MS Datasets 1 and 2

Frozen crosslinked cell pellets (600 mg wet cell mass) were resuspended in lysis buffer A (50 mM KCl, 25 mM HEPES, pH 7.3, 2.5 mM NaCl, 1 mM DTT, 0.625 mM MgCl_2_, 2.5% glycerol and 1% protease inhibitor) to 150 mg/ml and incubated with 0.3 mg/ml lysozyme for 30 min at 37°C. Immediately before sonication, 1 ml of lysis buffer B was added to a final concentration of 83.5 mM KCl, 42 mM HEPES, 4.2 mM NaCl, 1 mM DTT, 1.2 mM MgCl_2_, 4.2% glycerol and 1.5% protease inhibitor, and 2 µl benzonase was added to a concentration of 250 units/ml. Lysis by sonication was performed on ice using a Qsonica microtip probe (3.2 mm) for 30 seconds, 1 second on/1 second off with amplitude 12-24% on a Branson sonifier 250. The sample was kept on ice during sonication. After the last round, 2 ml lysis buffer B and additional DTT were added (final concentration: 100 mM KCl, 50 mM HEPES, 5 mM NaCl, 3 mM DTT, 1.5 mM MgCl_2_, 5% glycerol and 1.75% protease inhibitor) and the lysate was left to incubate for 30 min on ice. The lysate was clarified by centrifugation for 30 min at 20,000 x g and 4°C.

The supernatant was removed and the proteins were precipitated by chloroform/methanol precipitation (Wessel & Flügge, 1984), as material to produce Dataset 1. In parallel, the cell debris was washed with PBS and resuspended in 6 M guanidine hydrochloride with 50 mM Tris-HCl (pH 8) as before. The proteins were then precipitated with the chloroform/methanol precipitation, as material for Dataset 2. The samples for both datasets were processed separately but identically.

The precipitated pellets were processed as described in the proteomics section and peptides were cleaned up and stored on C18 HyperSEP cartridges at −80°C until use (Thermo Scientific).

As a first dimension of fractionation and crosslinked peptide enrichment, peptides were separated by strong cation exchange (SCX). Peptides were eluted from the C18 HyperSEP cartridges with 80% ACN, 0.1% TFA. Eluted peptides were dried in a vacuum concentrator and resuspended to a concentration of approximately 1.25 µg/µl in SCX buffer A (30% ACN, 10 mM KH_2_PO_4_). 400 µg were injected in SCX buffer A onto a PolySulfoethyl A SCX column (100 × 2.1 mm, 300 Å, 3 µm) with a guard column of identical stationary phase (10 × 2.0 mm), (PolyLC, Columbia, MD, USA) mounted on an Äkta pure system (Cytiva, Chicago, IL, USA) running at 0.2 ml/min at 21°C. After isocratic elution, a ‘step’ elution of 3.5% buffer B (30% ACN, 10 mM KH_2_PO_4_, 1M KCl) for 10 min eluted peptides that were discarded. Peptides were then eluted with increasing Buffer B and 200 µl fractions were collected. The elution was a series of linear gradients with the following targets: 3.5% at 0 min, 11% B at 11.5 min, 12.7% at 14 min, 14.5% at 15 min, 16.3% at 16 min, 18.8% at 17 min, 23.3% at 18 min, 30.3% at 19 min, 40.0% at 20 min, 70% at 21 min. Due to the limited amount of peptides that can be loaded on this column this process was repeated 6 times and the corresponding fractions were pooled to get enough material per fraction. In all, 24 fractions were carried forward for further processing. They were desalted using C18 StageTips, eluted, dried and stored at −80°C.

For a second dimension of fractionation and crosslinked peptide enrichment we separated each SCX fraction by size exclusion chromatography. Desalted peptides were resuspended in 25 µl 30% (v/v) ACN and 0.1% (v/v) TFA and treated for 1 min in a sonication bath. They were fractionated using a Superdex 30 Increase 10/300 GL column (GE Healthcare) with a flow rate of 10 µl/min using mobile phase 30% (v/v) ACN, 0.1% (v/v) TFA. 6 × 50 µl fractions at elution volumes between 1.1 ml and 1.4 ml were collected and dried in a vacuum concentrator.

Samples for analysis were resuspended in 0.1% v/v formic acid, 3.2% v/v acetonitrile. LC-MS/MS analysis was conducted in duplicate for SEC fractions, performed on a Q Exactive HF Orbitrap LC-MS/MS (Thermo Fisher Scientific, Germany) coupled on-line with an Ultimate 3000 RSLCnano system (Dionex, Thermo Fisher Scientific, Germany). The sample was separated and ionized by a 50 cm EASY-Spray column (Thermo Fisher Scientific). Mobile phase A consisted of 0.1% (v/v) formic acid and mobile phase B of 80% v/v acetonitrile with 0.1% v/v formic acid. LC-MS was performed at a flow rate of 0.3 μl/min. Gradients were optimized for each chromatographic fraction from offline fractionation ranging from 2% mobile phase B to 45% mobile phase B over 87 min, followed by a linear increase to 55% over 5.5 min, then an increase to 95% over 2.5 min. The MS data were acquired in data-dependent mode using the top-speed setting with a 2.5 second cycle time. For every cycle, the full scan mass spectrum was recorded in profile mode in the Orbitrap at a resolution of 120,000 in the range of 400 to 1,450 m/z. Normalized AGC = 3e6; Maximum injection time = 50 ms; Dynamic exclusion = 30 s; In-source CID = 15.0 eV. For MS2, ions with a precursor charge state between 3+ and 6+; Normalized AGC target = 5e4; Maximum injection time = 120 ms; Loop count = 10. Fragmentation was done with stepped-HCD collision energies 18, 24 and 30% and spectra were recorded with a resolution of 60,000 with the Orbitrap.

A recalibration of the precursor m/z was conducted based on high-confidence (<1% FDR) linear peptide identifications. The recalibrated peak lists were searched against the sequences and the reversed sequences (as decoys) of crosslinked peptides using the Xi software suite (version 1.7.6.4) (https://github.com/Rappsilber-Laboratory/xiSEARCH) for identification (Mendes *et al*, 2019). The following parameters were applied for the search: MS1 accuracy = 2 ppm; MS2 accuracy = 5 ppm; Missing Mono-Isotopic peaks = 2; enzyme = trypsin (with full tryptic specificity) allowing up to two missed cleavages; crosslinker = DSSO (with reaction specificity for lysine, serine, threonine, tyrosine and protein N termini); Noncovalent interactions = True; Maximum number of modifications per peptide = 1; Fixed modifications = Propionamide on cysteine; variable modifications = oxidation on methionine, methylation on glutamic Acid, deamidation of asparagine (only when followed by glycine in the sequence), hydrolyzed/aminolyzed DSSO from reaction with ammonia or water on a free crosslinker end. For DSSO, additional loss masses for crosslinker-containing ions were defined accounting for its cleavability (“A” 54.01056 Da, “S” 103.99320 Da, “T” 85.98264 Da). The database used was all proteins identified in each sample with an iBAQ > 1e6 (1716 proteins for Dataset 1, 1726 proteins for Dataset 2).

Prior to FDR estimation, matches were filtered for those with at least 4 matched fragments per peptide, for crosslinking to lysines or N-termini, and for having cleaved DSSO signature doublet peaks representing each matched peptide. The candidates were filtered to 2% FDR on protein pair level using xiFDR version 2.1.5.5 (https://github.com/Rappsilber-Laboratory/xiFDR) (Fischer & Rappsilber, 2017).

### Crosslinking MS Dataset 3

Frozen crosslinked cell pellets (600 mg wet cell mass) were used in dataset 3 preparation. Lysis was performed the same as for cells used for Datasets 1 and 2. The supernatant was further separated to simplify the crosslinked proteome to aid analysis. All steps were performed at 4°C. The lysate was clarified by centrifugation for 30 min at 20,000 x g. Soluble and insoluble proteome were separated by ultracentrifugation in a Beckman Coulter 70Ti fixed angle rotor at 38,000 rpm (100,000 x g) for one hour. The pellet was retained for digestion and crosslinking MS analysis. The supernatant was concentrated to 10% of the initial volume using a 100 kDa cutoff Amicon filter (Merck Millipore).

For lysate separation by size exclusion chromatography, 100 µl of concentrated lysate was loaded onto a Biosep SEC-S4000 (7.8 × 600) size exclusion column on an ÄKTA Pure (GE) Protein Purification System pre-equilibrated with running buffer (5% glycerol, 100 mM KCl, 50 mM HEPES, 5 mM NaCl, 1.5 mM MgCl_2_) and separated at 0.2 ml/min. 50 × 200 µl fractions were collected at elution volumes 10 ml (end of the void volume) to 20 ml. The fractions were pooled into 8 pools. The 8 protein pools were pelleted by acetone precipitation.

The 8 pools from protein SEC and the pellet from the ultracentrifugation step were digested as for Datasets 1 and 2 and stored on HyperSEP C18 SPE solid phase columns at −80°C prior to peptide fractionation. SCX plus subsequent SEC fractionation was performed for each pool of peptides as described for Datasets 1 and 2. Whenever amounts were insufficient, SCX fractions were pooled to have at least 20 µg prior to separation by SEC.

Samples were resuspended in 0.1% v/v formic acid, 3.2% v/v acetonitrile. LC-MS/MS analysis was conducted in duplicate for SEC and SCX fractions, performed on an Orbitrap Fusion Lumos Tribrid mass spectrometer (Thermo Fisher Scientific, Germany) coupled on-line with an Ultimate 3000 RSLCnano system (Dionex, Thermo Fisher Scientific, Germany). The sample was separated and ionized by a 50 cm EASY-Spray column (Thermo Fisher Scientific). Mobile phase A consisted of 0.1% (v/v) formic acid and mobile phase B of 80% v/v acetonitrile with 0.1% v/v formic acid. LC-MS was performed at a flowrate of 0.3 μl/min. Gradients were optimized for each chromatographic fraction from offline fractionation ranging from 2% mobile phase B to 45% mobile phase B over 100 min, followed by a linear increase to 55% over 5.5 min, then an increase to 95% over 2.5 min. The MS data were acquired in data-dependent mode using the top-speed setting with a 2.5 second cycle time. For every cycle, the full scan mass spectrum was recorded in the Orbitrap at a resolution of 120,000 in the range of 400 to 1,450 m/z. Normalized AGC = 250%, Maximum injection time = 50 ms, Dynamic exclusion = 60 s. For MS2, ions with a precursor charge state between 4+ and 7+ were selected with highest priority and 3+ were fragmented with any cycle time remaining. Normalized AGC target = 200%, Maximum injection time = 118 ms. Fragmentation was done with stepped-HCD collision energies 18, 24 and 30 % and spectra were recorded with 60,000 resolution with the Orbitrap.

Spectra recalibration, database search with xiSEARCH, and FDR thresholding with xiFDR was performed the same as for Dataset 1.

### CoFrac-MS

Cofractionation experiments were performed in triplicate on crosslinked and untreated cells as described in ‘Biomass production’. Lysis of cells was performed the same as described for crosslinking MS dataset 3 with lysate separated by size exclusion chromatography. 50 × 200 µl fractions were collected at elution volumes 10.5 ml to 20.5 ml. Proteins were pelleted by acetone precipitation. High-molecular weight range protein standards (Cytiva) were used to calibrate the elution profiles.

Protein digestion and peptide cleanup was performed as described above. 10% of each sample (by volume) was acquired on a Q Exactive HF Orbitrap Mass Spectrometer (Thermo Fisher Scientific, San Jose, USA) connected to an Ultimate 3000 UHPLC system (Dionex, Thermo Fisher Scientific, Germany). Settings were as described in ‘Proteomics for protein abundance estimation’.

Mass spectrometry raw data were processed using MaxQuant 1.6.12.0 under default settings with minor changes: two allowed missed cleavages; variable modifications per peptide: oxidation on Met, acetylation on protein N-terminal peptides, and for the crosslinked samples additionally DSSO-OH and DSSO-NH on lysines and N-termini. Carbamidoethylation on Cys was set as fixed modification. The database used covered all 4,191 proteins listed for *B. subtilis* 168 in UniProt (Reviewed Swiss-Prot). The ‘matching between runs’ feature was disabled. Protein quantification was done using the iBAQ approach. Proteins identified-by-site only, decoys and contaminants were discarded from the data.

Co-elution data was plotted using the seaborn 0.10.0 package (Waskom, 2021) with data normalized and smoothed with PCprophet v1.2 (Fossati *et al*, 2021). The raw elution profiles for three crosslinked and three untreated replicas are reported in **Supplementary Table S3**.

(Fossati *et al*, 2021) complex database for PCprophet and t(Fossati *et al*, 2021) GO enrichment (rf.txt, “POS”) of each replica and condition was assigned to all pairs making up the complexes. The highest score of each pair was used to compare the crosslinked with the non-crosslinked data (Table S4).

### CoFrac-MS analysis for candidate PPI generation with PCprophet

The MaxQuant output was filtered to remove ribosomal proteins, and the data was further filtered to proteins having at least three identified peptides and 9.5×10^6^ iBAQ in all three replicas of either the crosslinked or the untreated condition. CoFrac-MS analysis of both crosslinked and untreated conditions was performed with PCprophet v1.2 (Fossati *et al*, 2021) with standard settings. The complex database used by PCprophet was made up of interacting protein pairs downloaded from *SubtiW*iki, reduced to only those where both proteins are present in our filtered input data. We used the co-elution score prior to GO enrichment (rf.txt, “POS”) of each replica and condition, and assigned this value to all pairs making up the complexes. In order to only infer candidates within the SEC column resolving range, we only considered complexes of up to 10 members and with peak elution before 19.5 ml. The resulting protein pairs were filtered to a co-elution score of 0.8 or higher in at least 2 replicas of either the crosslinked or the untreated condition to retain only the highest confidence candidates. Each pairwise combination of proteins within the complexes was derived, yielding 667 protein-protein interactions submitted to AlphaFold-Multimer. The *Subti*Wiki repository was updated to include CoFrac-MS candidate interactions whenever these validated previous annotation, confirmed crosslinking MS interactions, or yielded high-confidence models.

### Calculating receiver operator characteristic (ROC) curves

366 non-homomeric interactions of the *Subti*Wiki database were considered as previously known interactions if they were predictable with the filtered input data of both conditions. To get a list of false interactions, the same number of random protein pairs with no homomeric or known interactions were selected. The PCprophet analysis was performed as described above, with these protein pairs serving as the complex database. During post-processing, co-elution scores in the range of 0.0 and 1.0 (0.1 steps) were used as cut-off values and, with the resulting list of protein pairs **(Table S4)**, true and false positive rates calculated to plot the ROC curves.

### Protein structure prediction

The full protein-protein interaction list from *Subti*Wiki (March 2022) (Pedreira *et al*, 2022) was filtered to remove interactions with homologous structures. Homology to the PDB was taken as a match by BLASTP (v. 2.9.0+) (Camacho *et al*, 2009) with Evalue < 1^-3^ and at least 30% sequence identity to a structure present in the PDB (database downloaded 16 Feb 2022). Paralogs mapping to the same PDB chains were retained. PPI candidate pairs from crosslinking MS, co-elution and *Subti*Wiki were further filtered to remove within-ribosome interactions. For experimentally-derived PPIs, protein pairs having homologs in the PDB were retained. Interactions were annotated as present in STRING version 11.5 (Szklarczyk *et al*, 2021) if their combined score exceeded 0.4.

2032 PPI candidate pairs were submitted to AlphaFold-Multimer v2.1.0 (Evans *et al*, 2022) (release November 2021, database downloaded 30 November 2021) and ran with full database size and the ‘is_prokaryote’ flag for MSA pairing switched off. Maximum structure template date was set to 1 November 2021. 5 models were predicted per run. A small fraction of runs ended in errors, and 1977 PPIs were modeled. Models were evaluated based on ipTM, pTM, predicted aligned error matrix and pLDDT score extracted from the runs. The top-ranking model by ipTM is used for the figures. For error control, 300 *B. subtilis* proteins from this dataset were predicted in complex with 300 random *E. coli* proteins and evaluated on the basis of ipTM score in relation to 10 subsamples of the 1977 PPIs predicted in the main dataset.

Accessible interaction volume for YugI and YabR AlphaFold models was computed using DisVis with a rotational search angle of 15° against the structure of the *B. subtilis* ribosome (PDB id 3j9w) (Sohmen *et al*, 2015). Crosslinking MS restraints were defined between 2.5 and 28 Å Cα-Cα. Crosslinks were mapped to structures using xiVIEW (www.xiview.org) and visualized using UCSF ChimeraX (Pettersen *et al*, 2021) and PyMol.

For trimer prediction, 33 trimers were submitted to AlphaFold-Multimer v2.2.1 (release June 2022, database downloaded 25 June 2022) based on dimers where the best model by ipTM had ipTM 0.65. AlphaFold v2.2.1 was run with full database size with 2 predictions with different random seeds per model. Maximum structure template date was set to 1 November 2021.

### Bacterial strains and plasmids

All strains are derived from the laboratory wild type strain *B. subtilis* 168. Deletion of the genes *yabR, yugI* and *pdhI* was achieved by transformation with PCR products constructed using oligonucleotides to amplify DNA fragments surrounding the respective genes and including an antibiotic resistance cassette as described (Guérout-Fleury *et al*, 1995). The same procedure was applied to fuse His-tags to the c-terminus of *yabR* and *yugI* and a FLAG-tag to *rocF*. The plasmid for overexpression of PdhI was constructed by amplifying *phdI* from chromosomal DNA and cloning the gene between a BamHI and a XbaI restriction site of the vector pBQ200 (Martin-Verstraete *et al*, 1994). For integration of the mutations into *pdhI*, overlapping primers carrying the desired mutations were used for the amplification of the fragment. The integration of or integration of the fragments into pBQ200n was carried out as described above.

### Genetic manipulation

Transformation of *E. coli* and the plasmid DNA extraction was performed using standard procedures (Sambrook *et al*, 1989). *B. subtilis* was transformed with plasmids, genomic DNA, or PCR products following a two-step protocol (Kunst & Rapoport, 1995). Transformants were selected on SP plates containing the appropriate antibiotics. Fusion polymerase, T4 DNA ligases and restriction enzymes were used according to the manufacturer. DNA fragments were purified *via* the QIAquick PCR purification kit (Qiagen, Hilden, Germany). DNA sequences were determined by Sanger sequencing. Chromosomal DNA from *B. subtilis* was isolated using the peqGOLDBacterial DNA Kit (Peqlab, Erlangen, Germany).

### Bacterial two-hybrid assay

To validate protein-protein interactions, a bacterial two-hybrid system based on an interaction-mediated reconstruction of the adenylate cyclase (CyaA) from *Bordetella pertussis* was used (Karimova *et al*, 1998). For this purpose, the two fragments of CyaA (T18 and T25) are fused to a bait and a prey protein. Interaction of these two proteins leads to functional complementation of CyaA and ultimately to the synthesis of cAMP. This is monitored by measuring the activity of a cAMP-CAP-dependent promoter of the *lac* operon that codes for ß-galactosidase in *E. coli*. The plasmids pUT18, pUT18C, p25N and pTK25 were used for the fusion of the proteins of interest to the T18 and T25 fragments of CyaA, respectively. The resulting plasmids are listed in Supplemental Table S8. The *E. coli* strain BTH101 was co-transformed with corresponding pairs of plasmids. Protein-protein interactions were visualized by plating the transformed strains on LB plates containing 100 µg/ml ampicillin, 50 µg/ml kanamycin, 40 µg/ml X-Gal (5-bromo-4-chloro-3-indolyl-ß-D-galactopyranoside), and 0.5 mM IPTG (isopropyl-ß-D-thiogalactopyranoside). The plates were incubated for 40 h at 30°C.

### Growth assays

To analyze the growth of *B. subtilis* mutant strains, the bacteria were cultivated in LB medium to inoculate precultures in MSSM minimal medium (Gundlach *et al*, 2017) containing glucose. The cultures were grown until the exponential growth phase was reached, harvested, resuspended in MSSM containing no carbon source and then the OD_600_ was adjusted to 0.2. This was used to inoculate the strains to an OD_600_ of 0.1 in a 96 well plate (Microtest Plate 96 Well, Sarstedt) in MSSM minimal medium containing the desired additions. Growth was measured using the Epoch 2 Microplate Spectrophotometer (BioTek Instruments) set to 37°C with linear shaking at 237 cpm (4 mm) for 24 h or 44 h. The OD_600_ was recorded every 10 min.

### Ribosome purification and Western blot of endogenously His-tagged YugI and YabR

For ribosome purification, wild type strain 168 and strains carrying His-tagged versions of YabR and YugI were grown in 1 l LB, the latter two containing additional 150 µg/ml spectinomycin (Sigma Aldrich), until an OD_600_ of 0.5. Cells of each strain were centrifuged for 15 min at 5000 x g and 4°C. Medium was discarded and the pellet cooled in an ice water bath. The ∼500mg pellet was dissolved in 2 ml Tico buffer (20 mM Hepes, 6 mM MgOAc, 30 mM KOAc, 2 mM DTT, pH 7.6) and lysozyme was added to a final concentration of 0.4 mg/ml. Cell lysis was achieved by freeze-thaw cycles on ice and completed with mild sonication on ice using a Qsonica microtip probe (3.2 mm) for 2x 15 seconds 1 second on/1 second off with amplitude 12-24% on a Branson sonifier 250. Genomic DNA was shredded by centrifugation in QIAshredder tubes (Qiagen) at 10,000 x g and 4°C for 2 min and digested by addition of RNAse-free DNAse I (Promega) for 10 min on ice. Lysates were clarified by centrifugation for 10 min at 10,000 x g at 4°C.

Ribosomes were separated from equal optical density units loaded onto 10-40% (w/v) sucrose gradients and centrifuged for 4 h at 32,000 rpm in a Sw-40 Ti rotor (Beckmann Coulter) at 4°C. Sucrose gradients were fractionated with a GradientStation (BioComp) monitoring A_260_. Protein was isolated from fractions of interest via ethanol precipitation. 40% of the protein material of each fraction was analyzed by Western blot using an anti-his antibody (Penta-His antibody, Qiagen #34660). Detection was performed with a secondary antibody conjugated with horseradish peroxidase (Anti-Mouse IgG Peroxidase antibody, A3682, Sigma Aldrich). All three blots were developed for 9.2 s after adding SuperSignal™ West Femto Maximum Sensitivity Substrate (Thermo Scientific).

### Integration of PPIs into *Subti*Wiki

The *Subti*Wiki repository was updated to include crosslinking MS interactions, excluding those within the ribosome and those involving the highly abundant ribosomal proteins (L7/bL12, RplL; L1/uL1, RplA; S3/uS3, RpsC), the elongation factors (Ef-Tu/Tuf and Ef-G/FusA), and the RNA chaperones (CspC, CspB). The *Subti*Wiki repository was updated to include CoFrac-MS candidate interactions whenever these validated previous annotation, confirmed crosslinking MS interactions, or yielded high-confidence models. The interactions can be assessed on the corresponding gene pages where they are shown in a graphical display. A click on the green line connecting two interaction partners gives a link to the relevant publications. Moreover, the PPIs are shown in the Interaction browser, an interactive network presentation.

High-confidence AlphaFold-Multimer predictions of the 153 binary (ipTM > 0.85) and 14 trimeric complexes (ipTM > 0.8) have been integrated in the Structure viewer carousel of *Subti*Wiki. To facilitate access to the predicted complex structures, a link to a complete list of all involved proteins is provided in the sidebar under “Special pages” (http://subtiwiki.uni-goettingen.de/v4/wiki?title=Predicted%20Complexes).

## Acknowledgements

We thank Dr. Panagiotis Kastritis and Dr. Steven Johnson for critical reading of the manuscript. We are grateful to Lily Rose for the help with some two-hybrid analyses. This research was funded by the Deutsche Forschungsgemeinschaft (DFG, German Research Foundation) under Germany’s Excellence Strategy – EXC 2008 – 390540038 – UniSysCat and project 426290502 and, in part, by the Wellcome Trust [Grant number 203149]. For the purpose of open access, the authors have applied a CC BY public copyright license to any Author Accepted Manuscript version arising from this submission.

## Author contributions

F.J.O’R. and J.R. initiated and designed the project. F.J.OR. and J.R. supervised and coordinated proteomic, CoFrac-MS and crosslinking MS experiments. C.F. and K.C. performed CoFrac-MS experiments. C.F. analyzed CoFrac-MS experiments. C.F. performed and analyzed proteomic experiments. C.F. and K.C. performed crosslinking MS experiments. F.J.O’R., S.L. and L.F. analyzed crosslinking MS results. A.G. performed and analyzed structural modeling. J.S. and R.B designed 2-hybrid experiments. R.B. performed and analyzed 2-hybrid and growth assays. C.F. performed sucrose gradient analysis. C.E. worked on data visualization and *Subti*Wiki integration. J.S. supervised and designed 2-hybrid, growth assays and *Subti*Wiki integration. J.R., F.J.O’R., A.G. and J.S. wrote the first version of the manuscript and all authors worked on manuscript completion and revision.

## Data Availability

Crosslinking MS data is deposited in JPOST and ProteomeXchange with identifiers JPST001796 and PXD035508 (Dataset 1), JPST001797 and PXD035519 (Dataset 2), JPST001791 and PXD035362 (dataset 3). CoFrac-MS data is deposited in ProteomeXchange JPOST with accession JPST001864 and PXD035520. Proteomic data is deposited in ProteomeXchange JPOST with accession JPST001795 and PXD03552. Top-scoring models are available in ModelArchive with accession ma-rap-bacsu. Protein-protein interactions and top-scoring models are added to the *Subti*Wiki repository (http://subtiwiki.uni-goettingen.de/).

