## supplementary_information for "Protein complexes in cells by AI-assisted structural proteomics"

**Table S1**: Crosslinking MS results, datasets 1-3 combined.

**Table S2**: Protein identifications and abundances for whole *B. subtilis* proteome.

**Table S3**: Co-elution data. iBAQ of all identified proteins in the SEC fractions for the crosslinked and untreated cells, including all replicates.

**Table S4**: PC-prophet co-elution scores for binary PPIs and data used for ROC curve.

**Table S5**: Summary of the top (by ipTM) models from AlphaFold-multimer runs of heterodimers.

**Table S6:** Summary of AlphaFold-multimer results for heterotrimers.

**Table S7:** Growth curves.

**Table S8:** Constructed plasmids.

**Figure S1:** Crosslinking MS workflow and resulting network.

**Figure S2:** Crosslinks visualized on structures.

**Figure S3:** Characterization of YugI-ribosome and YabR-ribosome interactions.

**Figure S4:** Co-elution profiles of known complexes from crosslinked and untreated cells.

**Figure S5:** Elution profiles of uncharacterized proteins with their partners detected by crosslinking MS.

**Figure S6:** Co-fractionation MS workflow using PCprophet for identifying candidate PPIs.

**Figure S7:** Models of novel interactions.

**Figure S8:** Groups of proteins connected through binary interactions predicted with an ipTM >0.65.

**Figure S9:** YclNOPQ structure.

**Figure S10:** pLDDT and uncertainty/flexibility in YneR.

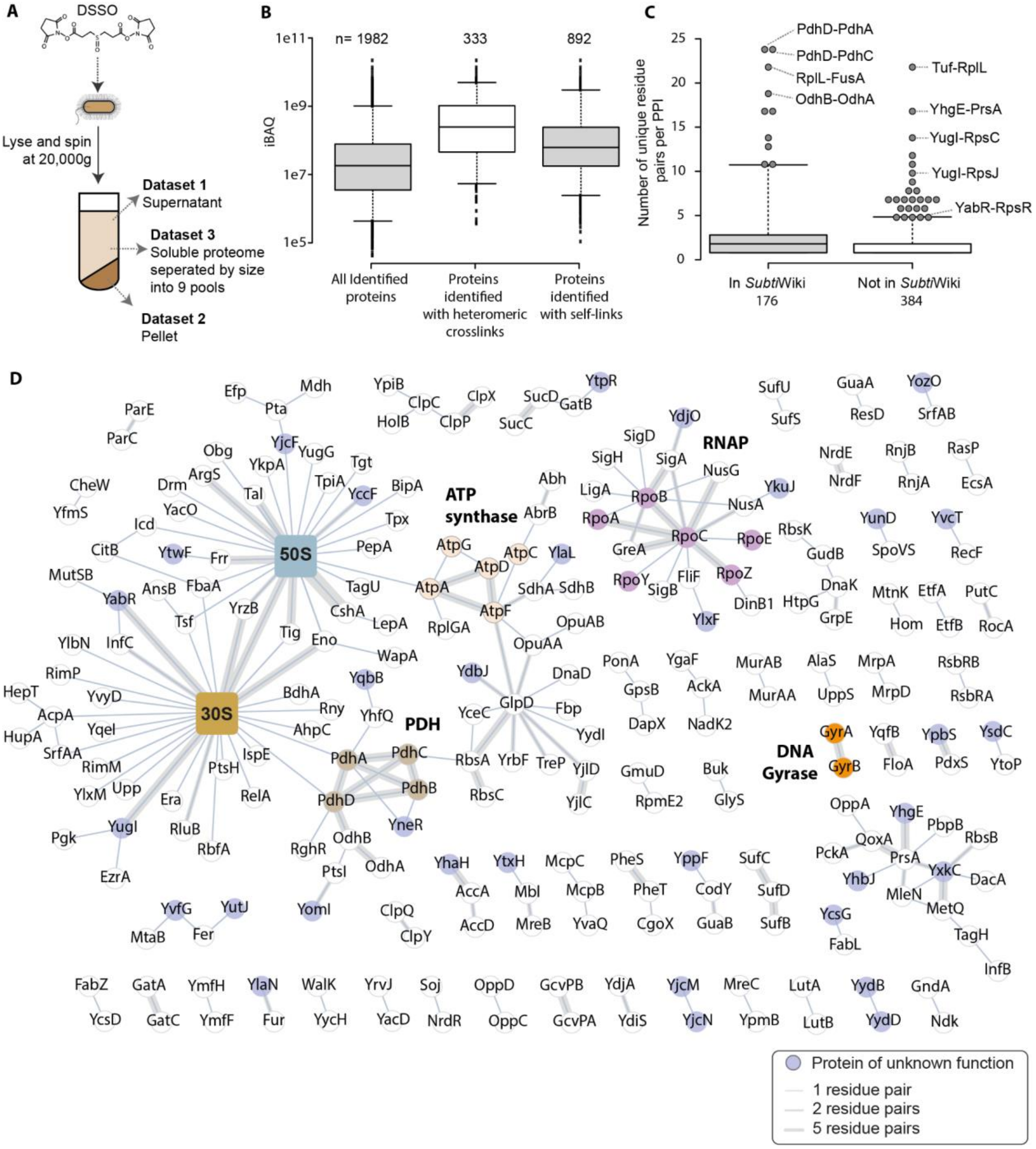


**Figure S1: Summary of crosslinking MS results.**

**A-** Cells were crosslinked with DSSO and the crosslinked proteome was separated into pellet and supernatant after lysis with a 20,000 x g centrifugation. Both were digested using Trypsin, the peptides were fractionated and Datasets 1 and 2 were acquired. The supernatant was also fractionated by size exclusion chromatography and high-molecular weight fractions were digested, the peptides fractionated and acquired as Dataset 3. **B-** The median intensity of proteins identified by shotgun proteomics is 1.8 x $10^7$, but for proteins with identified intra-protein or inter-protein crosslinks it is 6.2 x $10^7$ and 2.5 x $10^8$ respectively. **C-** Previously reported PPIs tend to have higher numbers of unique crosslinked residue pairs, suggesting higher abundance of these interactions/protein assemblies in the cell. **D-** PPIs identified at 2% PPI-level FDR (interactions to seven abundant and highly crosslinked proteins are removed for clarity). Previously uncharacterized proteins are shown in blue. Selected complexes are highlighted.

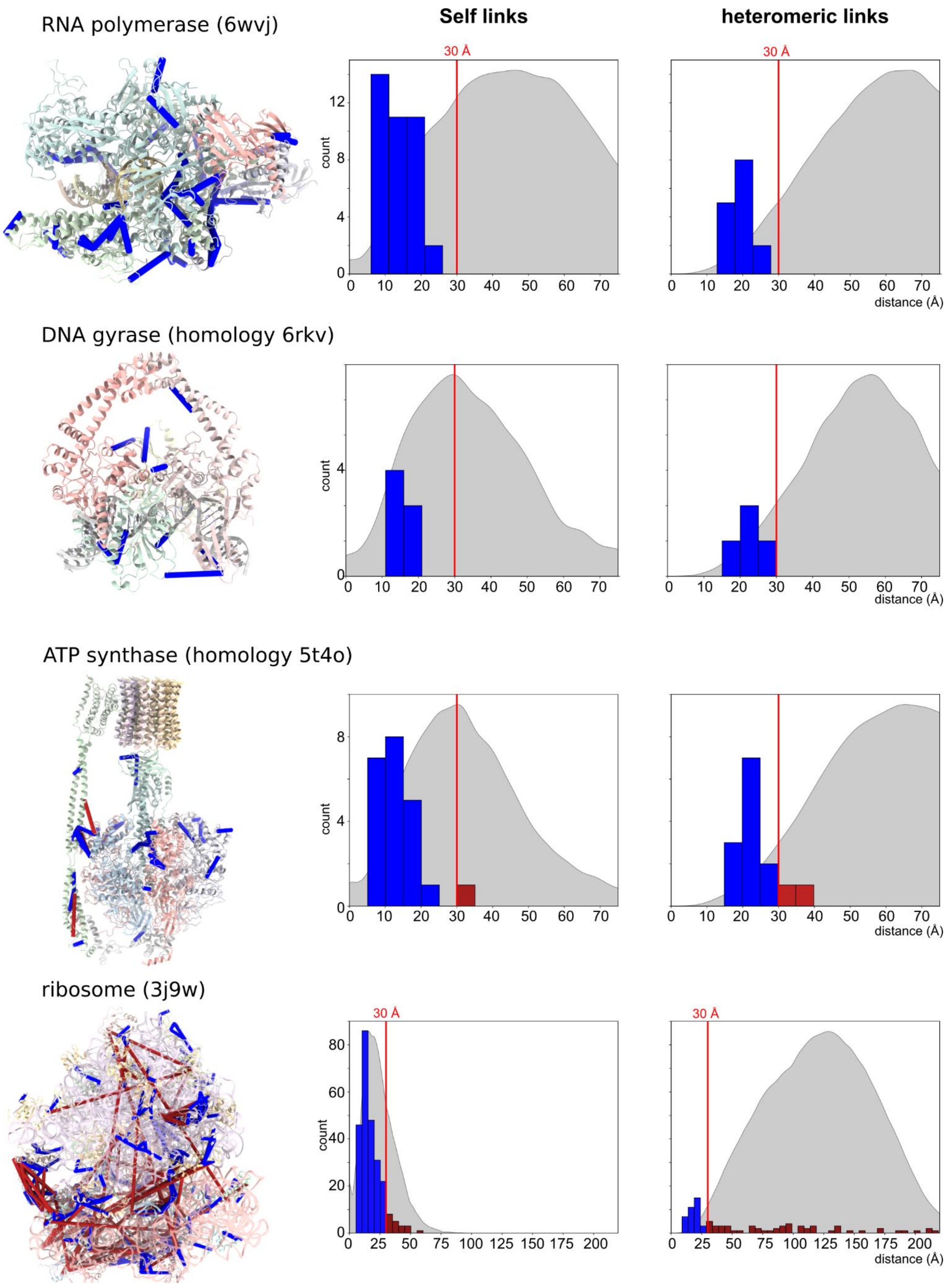

**Figure S2: crosslinks mapped onto structures of known complexes.**

Right: Mapping of crosslinking MS data on known complexes. Ribosome and RNA polymerase are represented by their experimental structures (Newing et al., 2020; Sohmen et al., 2015), while ATP synthase and DNA gyrase are modeled onto structures with high sequence identity (Sobti et al., 2016; Vanden Broeck et al., 2019). Satisfied crosslinks (<30 Å Cα-Cα) in blue, violated crosslinks in red. Crosslinks observed in ribosomes *in situ* include polysome contacts and contacts made in ribosome assembly intermediates. Left: distance distributions for self and heteromeric crosslinks in the structures on the left. The corresponding random distributions are overlaid in gray.

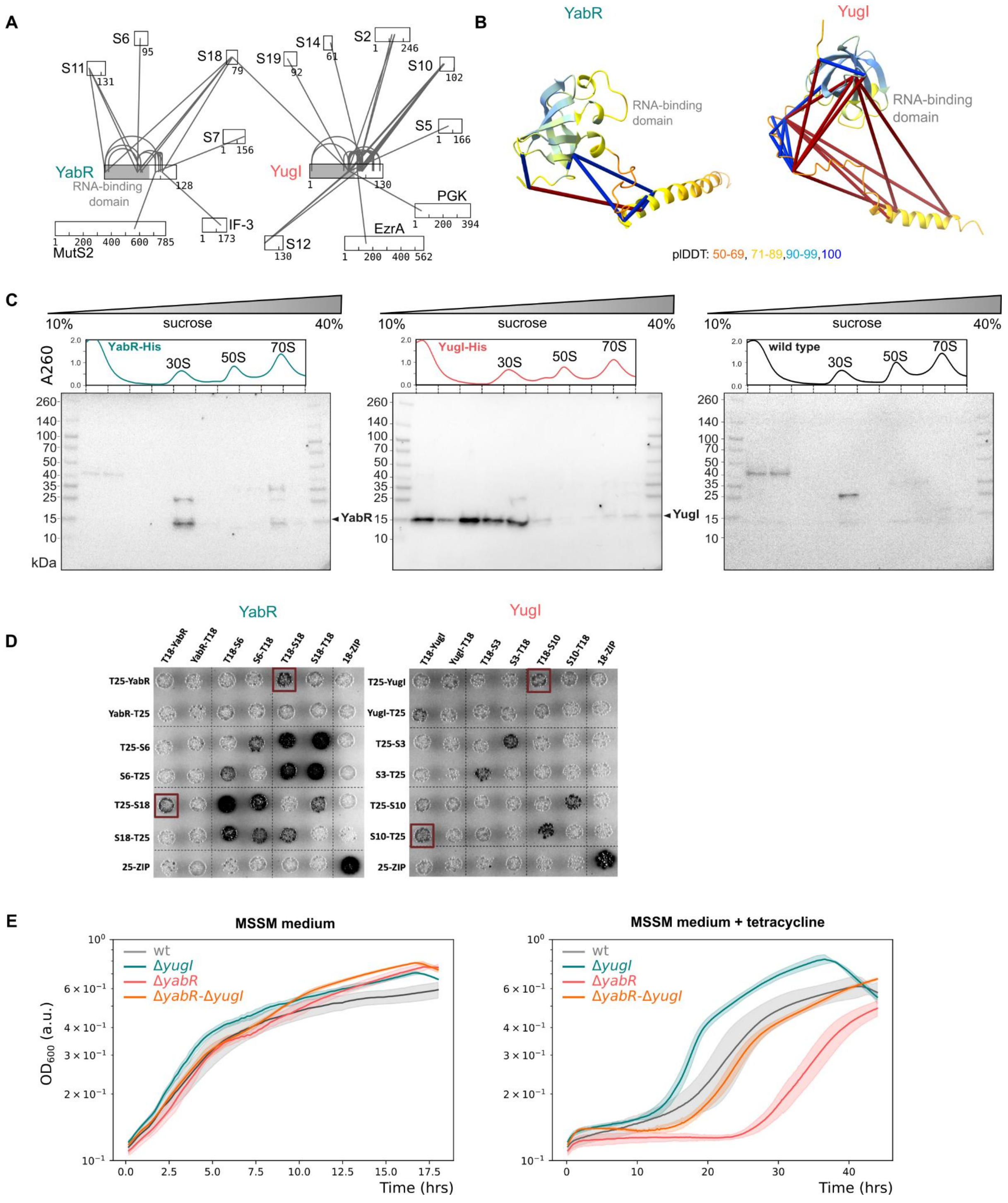


**Figure S3: Characterization of YugI and YabR ribosome interactions.**

**A-** Crosslinking MS network of YabR and YugI. Shaded area corresponds to S1-type RNA-binding domains in YabR and YugI. **B-** AlphaFold predictions of YabR and YugI colored by pLDDT with crosslinks shown. Crosslinking MS shows the high degree of flexibility of the C-terminal helices of the two proteins, which has low pLDDT score. Satisfied crosslinks (<30 Å Cα-Cα) in blue, violated crosslinks in red. **C-** Absorbance traces and western blot images relating from sucrose gradient separation of ribosomes from wild type strain 168 and strains carrying either YabR-His or YugI-His (**Fig. 1C**). **D-** Bacterial two hybrid experiment to identify interactions between YabR and the ribosomal proteins S6 and S18 as well as YugI and S3 and S10. All proteins of interest were fused to the T18 and T25 domains of the adenylate cyclase CyaA and interactions were tested in *E. coli* BTH101. Colonies turn dark as a result of protein interaction which enables adenylate cyclase activity and subsequently expression of the ß-galactosidase. A leucine zipper was used as a positive control. **E-** Resistance to tetracycline. Growth experiment of wild type (gray), Δ*yugI* (teal), Δ*yabR* (salmon) and Δ*yugI* Δ*yabR* (orange) in MSSM minimal medium with 0.1 mM KCl. The assay compares the growth under standard growth condition or after the addition of 5.2 mM tetracycline to the medium.

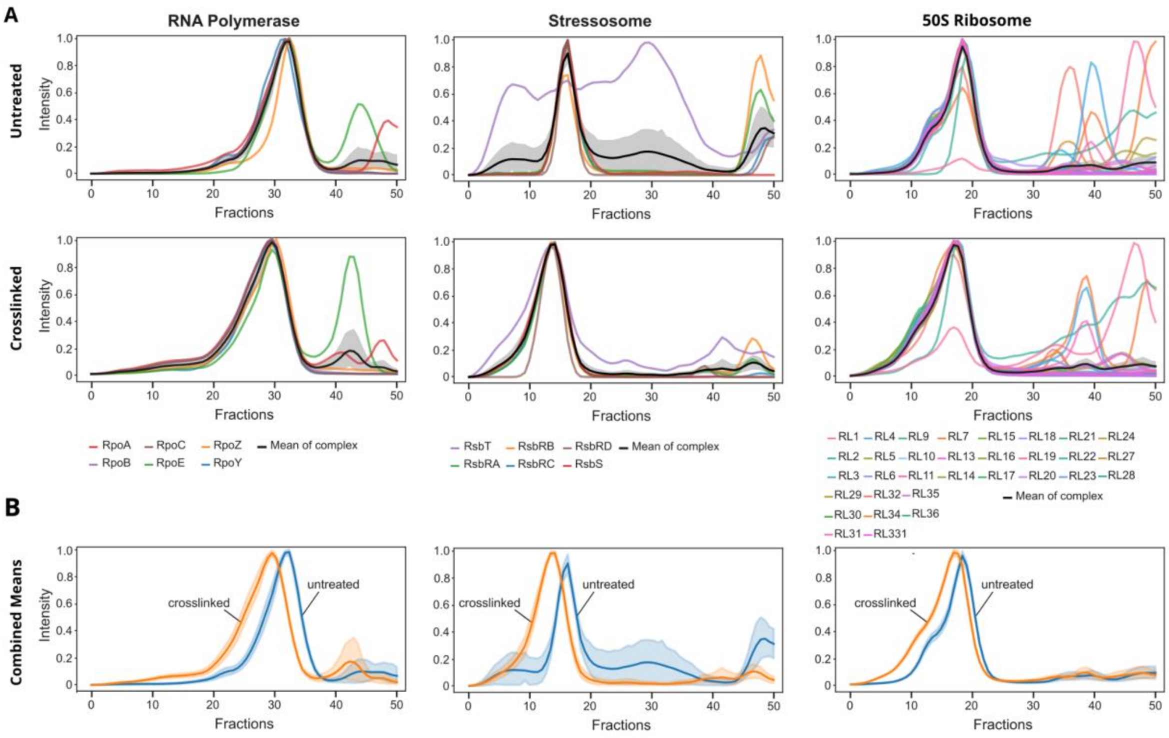


**Figure S4: Effects of crosslinking on elution behavior of known protein complexes.**

**A-** Elution profiles of core subunits of known complexes are shown as lines. The average normalized intensity across all subunits per fraction is in black, with the standard deviation in gray. Intensities are averaged across replicas. **B-** Comparison of the mean elution profile of the complexes from crosslinked and untreated cells.

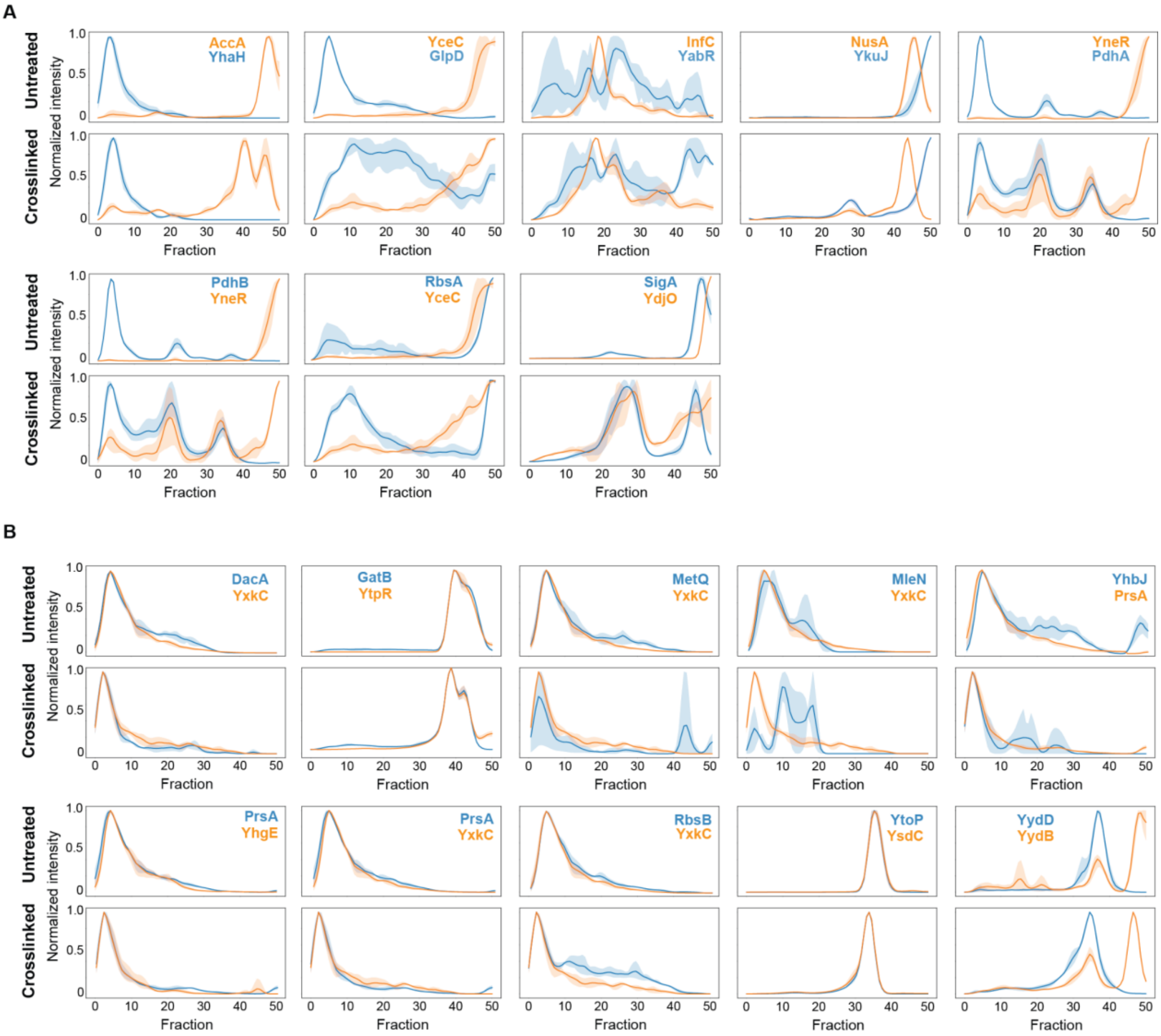


**Fig S5: Elution profiles of uncharacterized proteins with their partners detected by crosslinking MS.**

Manually annotated smoothed and normalized co-elution profiles of protein-protein interaction pairs involving uncharacterized proteins from untreated (upper panel) and crosslinked cells (lower panel). **A-** Protein pairs that show better co-elution behavior upon crosslinking, **B-** similar co-elution behavior.

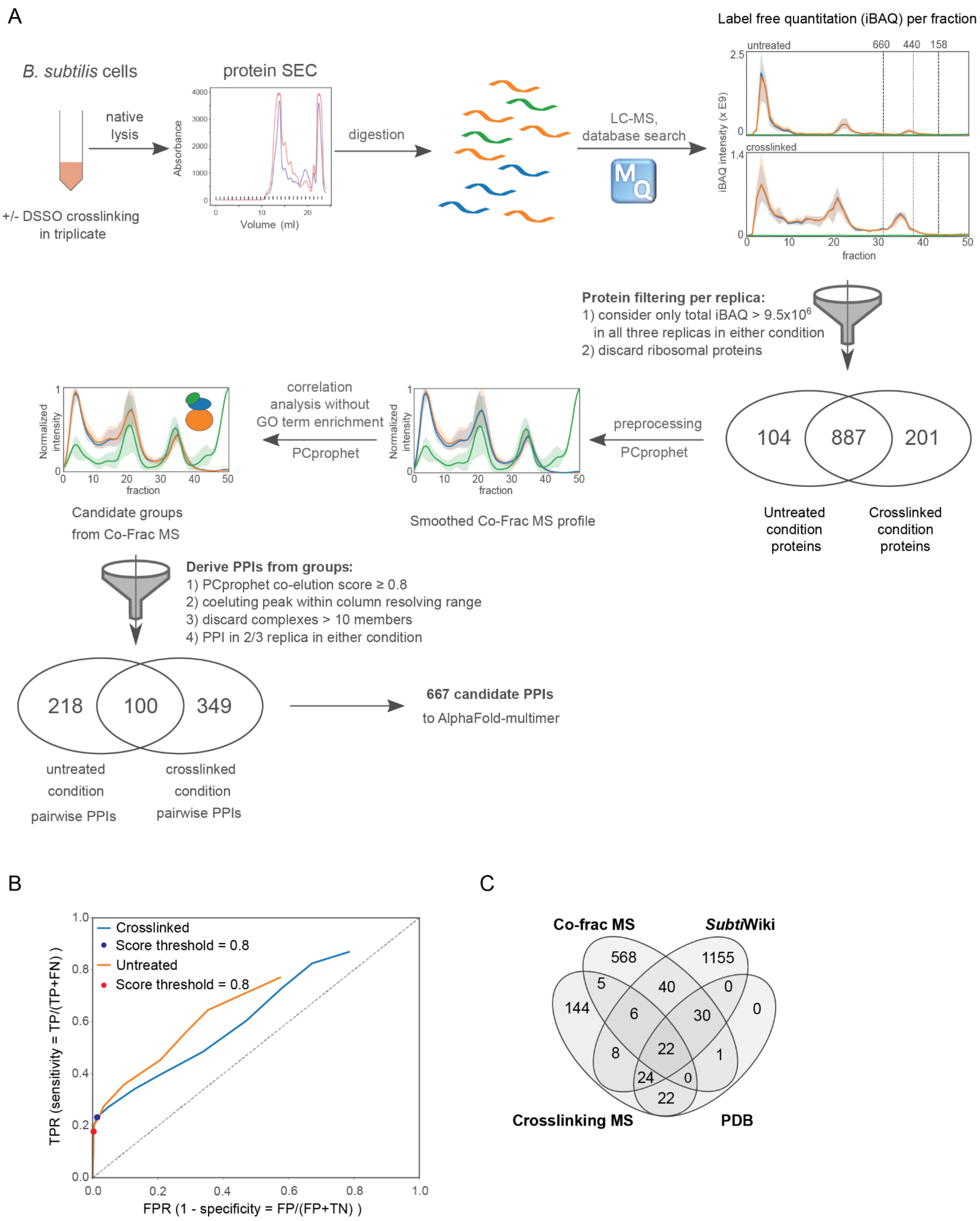


**Figure S6: Co-fractionation MS workflow using PCprophet for identifying candidate PPIs.**

**A-** CoFrac-MS workflow. **B-** Receiver operator characteristic (ROC) curves for the postprocessing of CoFrac-MS data for crosslinked and untreated cells. The threshold highlighted is chosen as the cutoff for submitting candidate PPIs to AlphaFold-Multimer, resulting in a true positive rate (TPR) of 0.178 and a false positive rate (FPR) 0.003 for untreated experiments, and a TPR of 0.232 and an FPR of 0.014 for crosslinked experiments.

**C-** Overlap of candidate PPI datasets along with previously known structures from the PDB (seq. identity > 30% and Evalue < $10^{-3}$). Note that *Subti*Wiki interactions were filtered to remove structurally characterized PPIs prior to candidate generation.

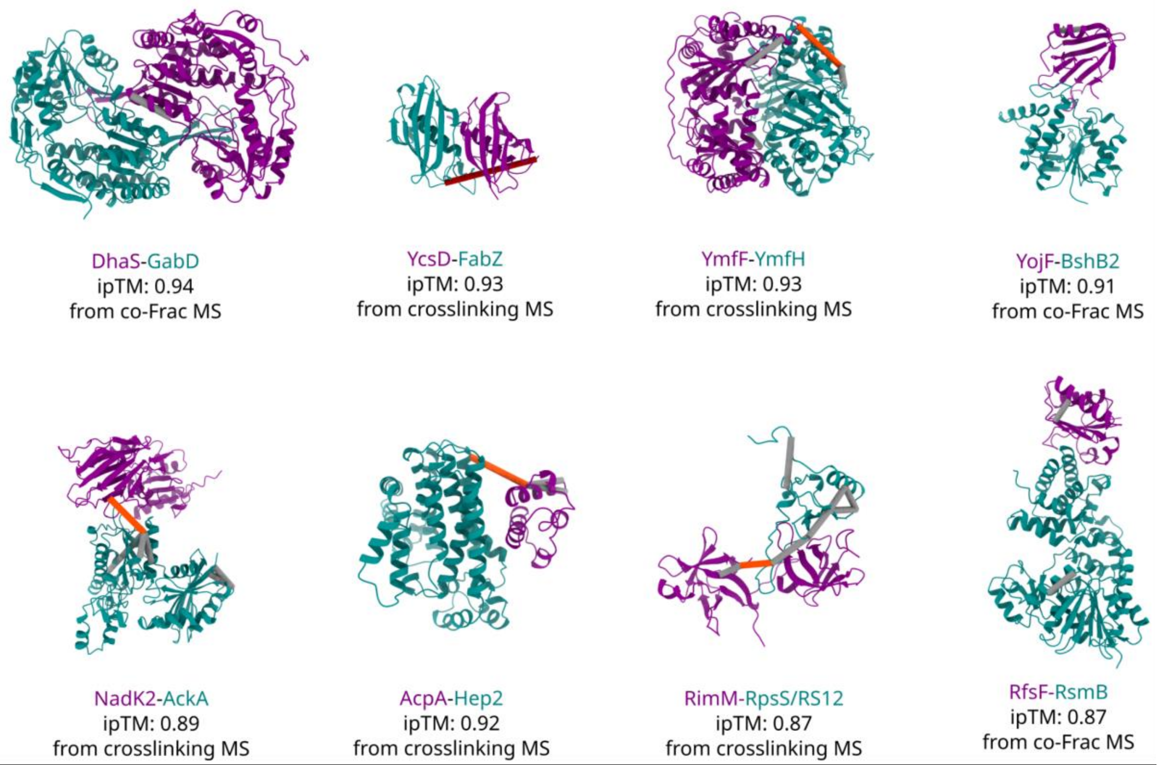


**Figure S7: Models of novel interactions.**

Models of high-confidence interactions (ipTM > 0.85) not previously annotated in *Subti*Wiki. Heteromeric crosslinks in orange, self crosslinks in gray.

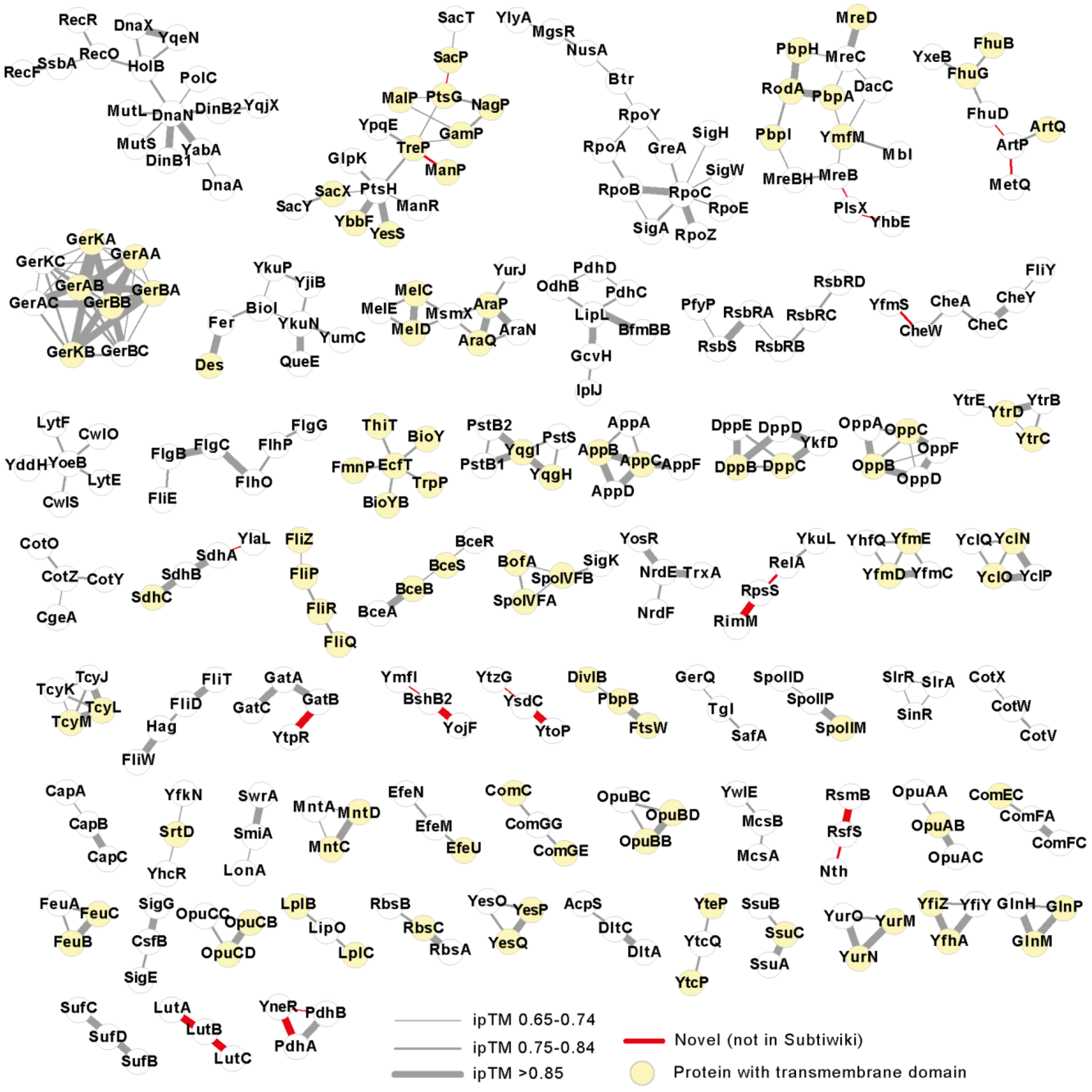


**Figure S8: Groups of proteins connected through dimers predicted with an ipTM >0.65.**

Groups containing only three members are predicted as 1:1:1 heterotrimers and shown in **Fig. 4**.

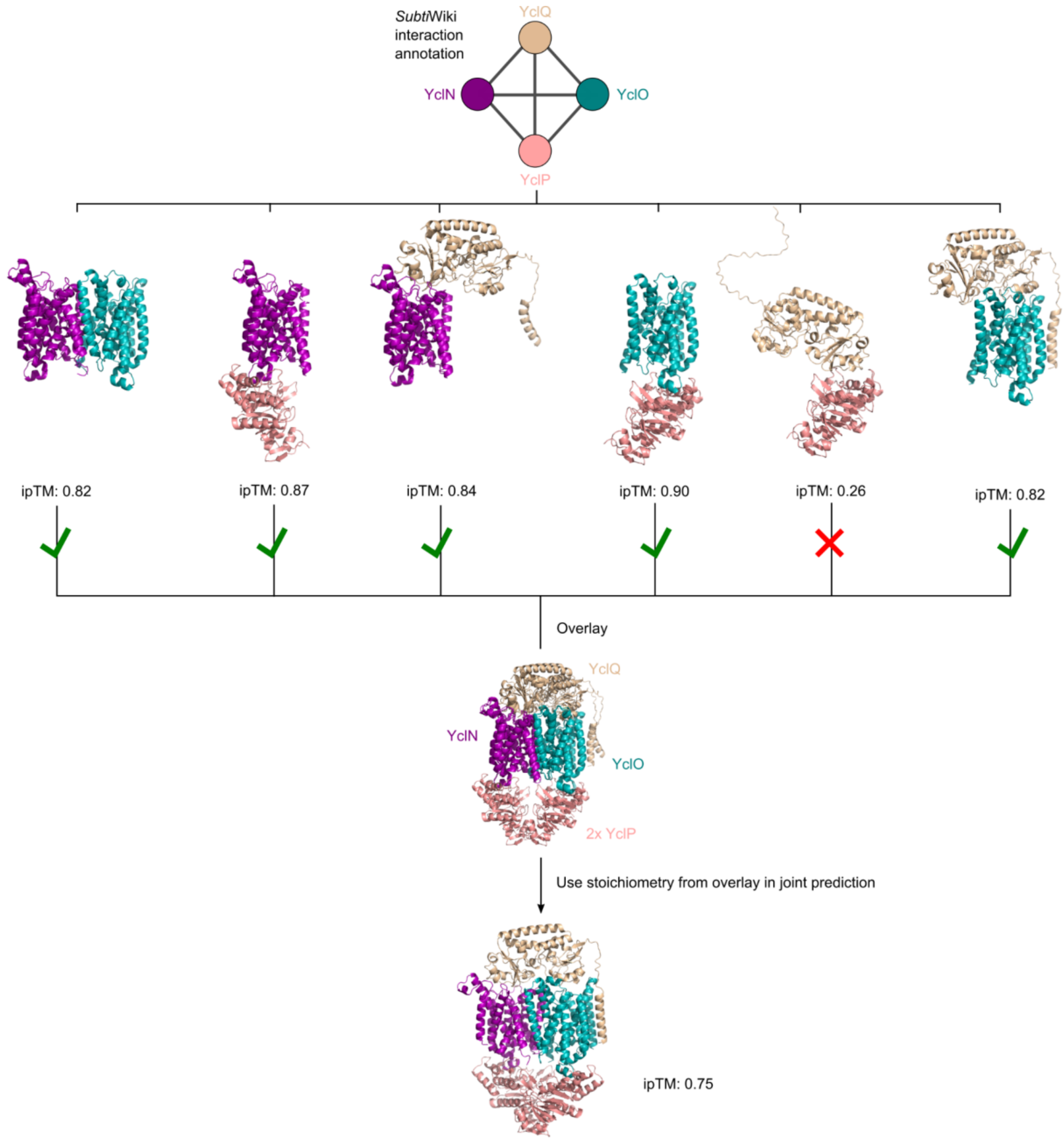


**Figure S9: YclN-YclO-YclP-YclQ assembly from pairwise predictions.**

The ABC transporter YclN-YclO-YclP-YclQ, involved in the transport of the siderophore petrobactin. The stoichiometry of this ABC transporter may be inferred by combining the predictions of its pairwise protein interactions. This information can be then used to select the stoichiometry used in predicting the whole complex.

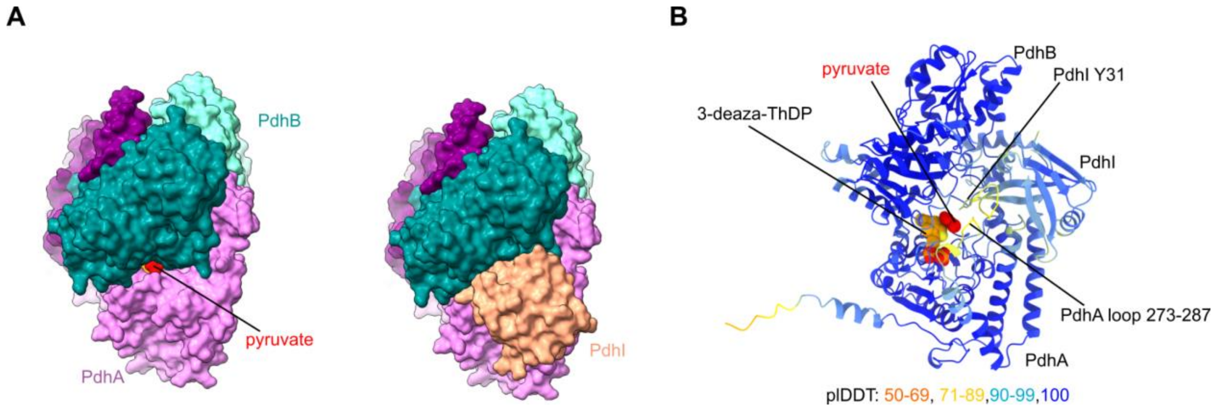


**Figure S10: The E1-PdhI interaction.**

**A-** Surface view of occlusion of the pyruvate binding site by PdhI. **B-** pLDDT residue coloring of the PdhI-PdhA-PdhB AlphaFold model highlighting flexibility and/or confidence in specific regions. Ligand positions as in pdb id 3dv0 (Pei et al., 2008).

Newing, T.P., Oakley, A.J., Miller, M., Dawson, C.J., Brown, S.H.J., Bouwer, J.C., Tolun, G., and Lewis, P.J. (2020). Molecular basis for RNA polymerase-dependent transcription complex recycling by the helicase-like motor protein HelD. Nat. Commun. *11*, 6420. .

Pei, X.Y., Titman, C.M., Frank, R.A.W., Leeper, F.J., and Luisi, B.F. (2008). Snapshots of catalysis in the E1 subunit of the pyruvate dehydrogenase multienzyme complex. Structure *16*, 1860–1872. .

Sobti, M., Smits, C., Wong, A.S., Ishmukhametov, R., Stock, D., Sandin, S., and Stewart, A.G. (2016). Cryo-EM structures of the autoinhibited E. coli ATP synthase in three rotational states. Elife *5*. https://doi.org/10.7554/eLife.21598.

Sohmen, D., Chiba, S., Shimokawa-Chiba, N., Innis, C.A., Berninghausen, O., Beckmann, R., Ito, K., and Wilson, D.N. (2015). Structure of the Bacillus subtilis 70S ribosome reveals the basis for species-specific stalling. Nat. Commun. *6*, 6941. .

Vanden Broeck, A., Lotz, C., Ortiz, J., and Lamour, V. (2019). Cryo-EM structure of the complete E. coli DNA gyrase nucleoprotein complex. Nat. Commun. *10*, 4935. .
